# SiglecF^high^ neutrophils in lung tumor tissues suppress local CD8 T cell responses and limit the efficacy of anti PD-L1 antibodies

**DOI:** 10.1101/2021.10.21.464997

**Authors:** Francesca Simoncello, Giulia Maria Piperno, Nicoletta Caronni, Tiziana Battini, Ambra Cappelletto, Silvio Bicciato, Federica Benvenuti

## Abstract

**Background:** Tumor infiltrating neutrophils generally correlates to worst prognosis and refractoriness to immunotherapy yet the complexity and significance of diverse subsets resident in tumor tissues has just begun to emerge. In lung tumors, a network of neutrophils states with graded protumorigenic properties is conserved between mouse and humans and include a subset of mature, long lived cells expressing the sialic-acid-binding protein SiglecF (SiglecF^high^ neu). The mechanism of recruitment of SiglecF^high^ neu into tumor tissues and the impact on local anti-tumor T cell responses and interference with immunotherapy is still elusive.

**Methods:** We used an immunogenic model of *Kras*^*G12D*^ Tp53 null adenocarcinoma of the lung to screen for factors inducing the recruitment of SiglecF^high^ neu, followed by gene editing to delete selected candidates. We analyzed frequencies and effector functions of endogenous CD8 T cell responses in controls and SiglecF^high^ neu depleted tumors by flow cytometry and functional assays. Tissues fluorescence and confocal imaging of lung sections was used to explore the relative distribution of neu and CD8 T cells. To establish the impact of SiglecF^high^ neu on anti-tumoral immune responses we treated cohort of animals with anti-PD-L1 antibodies to evaluate tumor growth in control conditions and under therapy.

**Results:** We found that tumor tissues express high levels of CXCL5, mapping to cancer cells. Upon deletion of chemokine expression by gene editing, the recruitment of SiglecF^high^ neu was almost entirely abrogated. In tumors depleted of SiglecF^high^ neu, the density of tumor specific endogenous CD8 T cells was 3-fold higher than in controls and showed significantly enhanced activation and effector functions. Importantly, checkpoint blockade with anti PD-L1 antibodies was ineffective in control tumors but showed a significant benefit in SiglecF^high^ neu depleted tumors.

**Conclusion:** This study demonstrates that SiglecF^high^ neu differentiating in lung tumor tissues inhibit local CD8 T cell responses and interfere with the success of checkpoint blockade. These data suggest that blocking selectively tissue resident neu may promote better responses to immunotherapy.

## Background

The impact of neutrophils (neu) on tumor progression has been extensively documented. Most experimental reports converge toward a pro-tumorigenic role of neu via promotion of cancer cell growth (1, 2), metastasis (3, 4, 5), stimulation of angiogenesis (6, 7) and modulation of T cell responses (8-10). Nevertheless, in some cancer types and tumor stages, neu possess anti-tumoral properties through direct killing of cancer cells and activation of T cell-dependent anti-tumor immunity (11, 12). In lung cancer, a high neu to lymphocyte ratio in peripheral blood is a well-established negative prognostic marker and neu density in tumor tissues has been associated to worst disease outcome (13-15). More recently, an abundant neu infiltrate has been associated to limited efficacy of checkpoint inhibitors in various cancers, including lung cancer (16-18) and neu density in tumor tissues was directly linked to decreased local antitumoral CD8 responses (19). Single cell sequencing and mass-cytometry has begun to unravel the diversity and complexity of neu subsets in tumor bearing hosts, underscoring the presence of different states with diverse tumor promoting potential and the importance to focus on neu infiltrating, non-ectopic, relevant tumor tissues (20-22). In particular, the structure of neu subsets in lung tumor tissues is well conserved between human lung adenocarcinomas and the orthotopic Kras^G12D/WT^; Tp53 (KP) mouse model (20, 23). KP lung tumors contain a dominant population of SiglecF expressing neu, representing aged neutrophils with high glycolytic activity that differentiate in situ (23-25). SiglecF expressing neu have been identified as well in other inflammatory conditions (26, 27), yet the exact tissue cues determining upregulation of SiglecF and the functional significance of its expression are only partially understood. In lung tumors, SiglecF^high^ neu were shown to produce reactive oxygen species and to promote cancer cells growth, however the impact of SiglecF^high^ neu accumulation of anti-cancer T cell responses and the interference of these cells with immunotherapy is still limited.

In this study, we identified tumor-derived CXCL5 as crucial to promote accumulation of SiglecF^high^ neu in lung tumor tissues. Genetic deletion of *Cxcl5* inhibited recruitment of SiglecF^high^ neu in lung tumors and promoted expansion and activation of tumor specific CD8 T cells. Importantly, depletion of SiglecF^high^ neu increased the sensitivity of KP tumors to PD-L1 checkpoint blockade.

## Material and Methods

### Mice

C57BL/6 and OT-I (C57BL/6-Tg(TcraTcrb)1100Mjb/J) mice were purchased from Envigo or Jackson Laboratories respectively. Animals were maintained in sterile isolators at the ICGEB animal Bio-experimentation facility (12h/12h light and dark cycle, 21 °C ± 2°C).

Sample size was determined based upon prior knowledge of the intragroup variation of tumor challenges by our laboratory and published studies ((24, 28) and was sufficient to detect meaningful biological differences with good reproducibility.

### Cell lines

The KP cells (LG1233) were generated from lung tumors of C57BL/6 KP mice (K-ras^LSLG12D/+^;p53^fl/fl^ mice) and kindly provided by Dr. Tyler Jacks (Massachusetts Institute of Technology, Cambridge, USA) (29). KP OVA cells were generated by transduced with the lentiviral vector Pdual-liOVAha-PuroR as described in (30). To generate KO^CXCL5^ cells, the KP OVA cells were transiently co-transfected with pSpCas9(BB)-(PX458) and pZac2.1-U6sgRNA-CMV-ZsGreen plasmids, carrying 5’-caccgCTGCCGCAGCATCTAGCTGA-3’ guide. The ZsGreen^+^ cells were sorted and single clones were tested for CXCL5 expression by ELISA (abcam ab100719).

To rescue CXCL5 expression in KP OVA KO^CXCL5^ cells were transduced with a lentivirus carrying the expression vector pLVX-IRES-G418-CXCL5 or empty vector pLVX-IRES-G418 as control. After antibiotic selection, CXCL5 expression was tested by ELISA.

All cell lines were maintained in DMEM media (containing 1g/L of glucose) supplemented with 10% fetal bovine serum (FBS, Euroclone) and Gentamicin (50μg/mL, Gibco) and routinely tested for mycoplasma contamination. The growth rate *in vitro* was assessed by crystal violet assay.

### Tissue preparation for flow cytometry

Lung tissues from control or tumor bearing mice were harvested after PBS lung circulatory perfusion, mechanically cut into small pieces and digested with 0,1 % Collagenase type 2 (265U/mL; Worthington) and DNase I (250U/mL; Thermo scientific) at 37°C for 30’. Cells were filtered using 70μm cell strainer (Corning), to obtain single cells suspension. Blood was collected through subclavian vein puncture, follow by red blood cell lysate. Spleen, lymph nodes (mediastinal and inguinal) were smashed and filtered with a 70 μm and 40 μm cell strainer respectively. Bone marrow was extracted from femur and tibia, flush to obtain a sing single cell suspension.

### Flow cytometry

For cell staining, FcR binding sites were blocked by using αCD16/CD32 and viability of cells was assessed by staining with LIVE/DEAD dye). The antibodies used for the experiments are listed in Supplementary Table 1. To analyze *Cxcl5* expression, CD45^+ or –^ populations from healthy or tumor bearing lungs were stained with CD45-A647 antibody and sorted. MHC-I-OVA pentamers (SIINFEKL/H-2Kb Pro5, Proimmune) were used to identify OVA specific CD8 T cells following manufacturer’s instruction. For intracellular detection of IFNγ, single cell suspensions were stimulated with OVA peptide (SIINFEKL) 2µM 37°C for 4 hrs in the presence of Golgi Stop (monensin, BD Biosciences). Upon extracellular staining, cells were fixed and permeabilized using Cytofix/Cytoperm solution (BD Biosciences) following manufacturer’s instructions.

To identify effector EOMES^-^Tbet^+^ CD8^+^ T cells, we performed a nuclear staining analyzed using Foxp3/transcription factor staining buffer set (ThermoFisher) following manufacturer’s instructions. To measure OVA expression, cells were fixed and permeabilized using Cytofix/Cytoperm solution (BD Biosciences) following manufacturer’s instructions, stained with rat-monoclonal αHA and then with αRAT-AF488. Where indicated, absolute cell count was analyzed by adding TrueCount Beads (Biolegend) to the samples following manufacturer’s instructions. Samples were acquired with FACS Celesta (BD Biosciences) and analyzed with FlowJo software (Tree Star, Inc.).

Flow cytometry experiments are based on objective measurements and blinding was not required.

### Real time PCR

RNA from total lungs or sorted cells was extracted using Trizol reagent (ThermoFisher Scientific), according to manufacturer’s instruction. cDNA was synthesized using SuperscriptII (ThermoFisher) and quantitative real-time PCR (qRT-PCR) was performed using SsoFast EvaGreen Supermix (Biorad) with specific primers: Gapdh For (AGAAGGTGGTGAAGCAGGCAT) Rev (CGAAGGTGGAAGAGTGGGAGT), Cxcl5 For (GCT GCC CCT TCC TCA GTC AT) Rev (CAC CGT AGG GCA CTG TGG AC). Gene expression profiling of inflammatory cytokines and receptors of normal and KP-OVA tumor bearing lungs was performed by custom RT^2^ Profiler PCR Array (Qiagen, cat. 330221) following manufacturer’s instructions.

### Immunohistochemistry

To assess tumor burden, lung tissues were harvested and fixed in formaldehyde 10% and paraffine embedded following standard procedure. Consecutive sections of 8µm were dewaxed and rehydrated and stained with the H&E using (Bio-Optica, Milano Spa). The area of tumor nodules was quantified manually over consecutive sections and averaged (3 sections/sample).

To identify neutrophils or proliferating cells within nodules, sections were treated with antigen-retrieval solution (Vector laboratories) for 20 min at 120°C. Slides were treated for 10 minutes in H_2_O_2_ and after blocking in 10% goat serum in 0.1% Tween20 for 30 minutes and incubated overnight at 4°C with specific antibody diluted in PBS 0,1%Tween20: anti-mouse Ly6G (1A8, BD Pharmingen, cat 551459), or anti-mouse Ki67 (D3B5, Cell Signaling, cat 12202s). Detection was performed using the ImmPRESS polymer detection system (Vector Laboratories), according to manufacturer’s Instructions. Automatic thresholding and measurements were performed using Ilastik or imageJ software, respectively. Images were acquired by Leica microscope. For tumor burden, neutrophil and proliferating cells measurements slides were scored blindly by two independent operators.

### Immunofluorescence staining

To identify SiglecF^high^ neu, tumor tissues were intratracheally perfused with 1% paraformaldehyde (PFA), fixed in 4% PFA and embedded in a frozen tissue matrix following standard procedure. Sections of 5µm were dried 10’ at RT, fixed with 4% PFA for 15’ and permeabilized for 15’ with PBS 0,5% Triton. After blocking in 5% mouse serum in PBS 1%BSA/0,1%NP-40 for 30’, slides were incubated overnight at 4°C with specific antibody diluted in PBS 1%BSA/0,1%NP-40: anti-mouse Ly6G-PE and anti-mouse SiglecF-BB515 (listed in Supplementary Table 1). Nucleus were labelled by Hoechst 15’ at RT. Images were acquired with LSM880 META reverse microscope.

To assess spatial distribution of neutrophils and CD8 T cells, tumor tissues were paraffine embedded. Sections of 8µm were dewaxed, rehydrated, treated with antigen-retrieval solution, and incubated overnight at 4°C with rat anti-mouse CD8 (4SM15, Invitrogen) and rabbit anti-mouse Ly6G (E6Z1T, Cell Signaling) antibodies diluted in PBS 0,1%Tween20 followed by α-rat 647 (Invitrogen) and α-rabbit AF488 (Invitrogen) nucleus were labelled by Hoechst.

Images were acquired with C1 Nikon reverse microscope. Automatic thresholding was performed by Ilastik. ImageJ software was used to quantify CD8 T cells and neu and to measure nodule’s area. Nodules having an area <0,09 mm^2^ were classified as small, the ones having an area ranging from 0,091 to 0,2 mm^2^ as medium and those with an area >0,2 mm^2^ as large nodules. Spatial distribution of neu and T cells was performed blindly on code-labeled slides.

### In vivo tumor challenge and blocking studies

To establish the adenocarcinoma tumor models C57BL/6 WT female mice at 8-10 wks of age, were intravenously injected with 7×10^4^ KP cells or with 2×10^5^ KP OVA variants (WT or KO^CXCL5^, KO^CXCL5 (lenti-CXCL5)^ or KO^CXCL5 (lenti-vec)^). Otherwise indicated, mice were sacrificed at initial (9 days) or at established stage of tumorigenesis (18 days). To deplete neutrophils *in vivo*, KP OVA bearing mice were intraperitoneally treated every 3 days, starting from the 6^th^ day to 12^th^ day, with 200 µg of anti-Ly6G antibody (InVivo Plus, clone 1A8, Bio X Cell) or isotype control (Rat IgG2a isotype control, clone 2A3, Bio X Cell) and sacrificed at day 13^th^.

In PD-L1 blockade experiments, mice were challenged with KP OVA WT or KO^CXCL5^ cells, intraperitoneally treated every 3 days with 200µg of αPD-L1 (InVivoMab, clone 10F.9G2, BioXcell) or isotype control (InVivoMab, rat IgG2b isotype control, clone LTF-2, BioXcell). Mice were sacrificed at day 18^th^.

For adoptive transfer of tumor specific T cells, mice were challenged with WT or KO^CXCL5^ cells. 9 days after 2×10^6^ OTI^+^CFSE^+^ T cells were intravenously injected and after 2 days mice were sacrificed to assess IFN-γ production in OTI T cells.

### Human gene expression data

Details on collection and processing of human lung cancer gene expression data, human neutrophil fraction analysis and survival analysis are indicated in supplementary methods.

### Statistic

Primary data were collected in Microsoft Excel and statistical analysis were performed by using Graphpad Prism 8 software. Values reported in figures are expressed as the standard error of the mean, unless otherwise indicated. For comparison between two or more groups with normally distributed datasets 2-tailed Student’s T test, multiple T test, one-way ANOVA or 2-way ANOVA were used as appropriate. For the comparison of matched groups, we used Wilcoxon test. The non-parametric Kruskal-Wallis test with Dunn’s multiple comparison was performed to compare 3 or more unmatched groups. p values > 0.05 were considered not significant, p values ≤ 0.05 were considered significant: * p≤0.05, ** p≤0.01, *** p ≤0.001, **** p ≤0.0001.

No exclusion criteria or data has been performed.

## Results

### Parental KP and immunogenic KP lung tumors are dominated by SiglecF^high^ neu

To corroborate sparse indications on the impact of neu infiltration for disease outcome, we first analyzed a large compendium of lung adenocarcinoma (LUAD) (Table 1), that integrates multiple RNAseq datasets. Stratification of patients based on low and high neu content using CIBERSORT (31) showed a clear correlation of high neu content with bad survival.

**Table 1.**
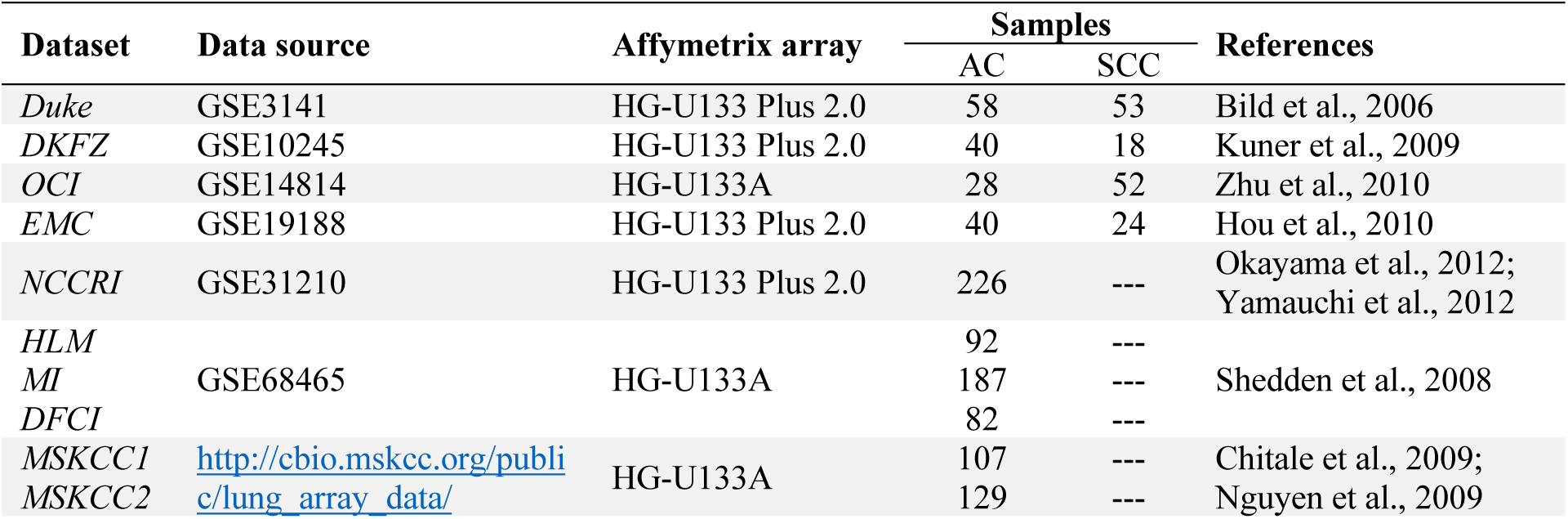
Datasets composing the lung cancer compendium (AC: adenocarcinoma; SCC: squamous cell carcinoma).

A second stratification method, based on expression of transcripts that define intratumoral neu (20), confirmed that tumor tissue neu are associated to worst outcome (Fig 1A). To study neu function, we next moved to the KP lung adenocarcinoma model that reflects features of the human disease and it is highly infiltrated by neu (20, 32). In parallel, we analyzed the recently generated OVA expressing immunogenic KP counterpart (KP-OVA), to allow tracking of tumor specific T cell responses, (30). KP-OVA lung tumors were dominated by a large proportion of CD11b^+^/Ly6G^+^ infiltrating cells, similarly to the non-immunogenic tumors (Fig 1B). Both KP and KP-OVA showed a slight decrease in CD4 and B cells, as compared to normal lungs and KP-OVA showed, as expected, an increment in the fraction of CD8 T cells (Fig 1B and S1A). To further phenotype tumor infiltrating neu, we analyzed expression of SiglecF, which was recently reported to mark a specific subpopulation of long-lived, mature lung tumors neu, distinct from other myeloid cells (23-25). Up to 70% of CD11b^+^/Ly6G^+^ cells in both KP and KP-OVA tumors expressed high SiglecF (SiglecF^high^ neu), whereas the majority of neu in normal lungs, and a minor proportion in tumors, were SiglecF low (SiglecF^low^ neu) (Fig 1C). A small fraction of SiglecF^high^ neu was found as well in the tumor draining lymph node (mediastinal, mLN). In contrast, we found no SiglecF^high^ neu in the blood, bone marrow, spleen or non-draining lymph nodes, indicating that this subset is not present systemically in tumor bearing animals, but it is restricted to lung tumor tissues and the connected lymph node (Fig 1D). We next used tissue imaging to visualize the distribution of neu within KP nodules.

**Figure 1.**
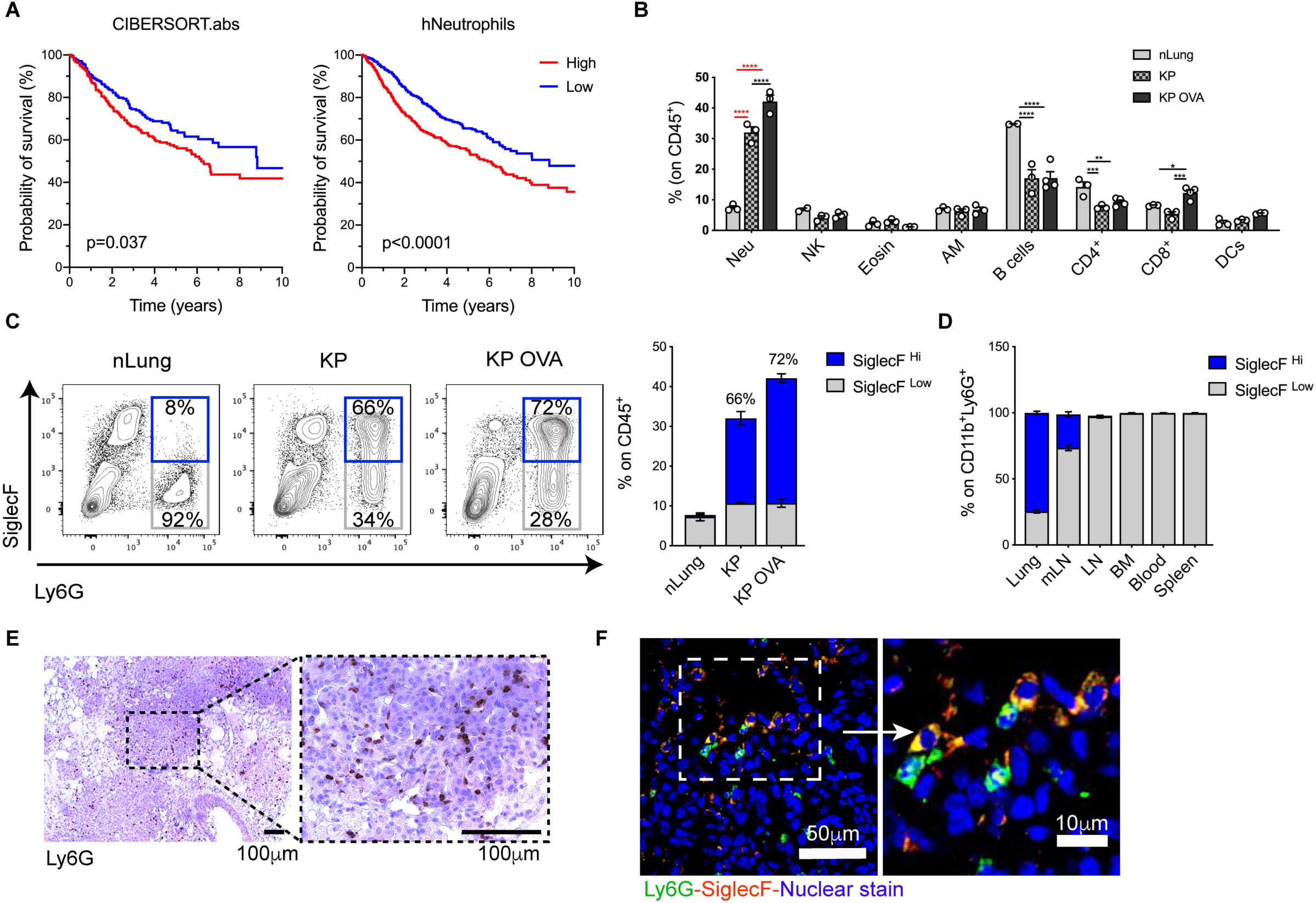
Characterization of neutrophils density in lung adenocarcinoma patients and in the KP model. **A)** Overall survival of LUAD patients with the highest and lowest neu proportions calculated with CIBERSORT (left) and the highest and lowest expression of infiltrating human neu (hNeutrophils, right) gene signature (Supplementary Table 1). **B-E)** Mice were challenged intravenously with KP, KP-OVA cells or PBS (nLung). The immune composition of lung tumor tissues was evaluated by flow cytometry in normal and established tumor bearing lungs. **B)** Relative abundance of each subset expressed as a fraction of CD45^+^ cells (neu=neutrophils, NK= Natural Killer cells, Eosin= Eosinophils, AM= Alveolar Macrophages, B cells= B lymphocytes, CD4^+^= CD4 T lymphocytes, CD8^+^= CD8 T lymphocytes, DCs= Dendritic Cells). Data represents mean±SEM of 2-4 mice each group. Significance was determined by two-way ANOVA with *p≤0.05; **p≤0.01; ***p≤0.001; ****p≤0.0001. **C)** Representative dot plot of SiglecF staining. Numbers in quadrant show percentages of SiglecF^high/low^ among CD11b^+^Ly6G^+^ cells. Bars show quantification of neu frequencies on CD45^+^ total lung cells and the fraction of SiglecF^high/low^ among neu (percentages above bars refer to SiglecF^high^ neu). Data are mean±SEM of 3-5 mice/group. **D)** Percentage of SiglecF^high^/^low^ neu within CD11b^+^Ly6G^+^ cells in the indicated organs isolated from KP OVA tumor bearing mice (mLN=mediastinal lymph node, LN=inguinal lymph node, BM=bone marrow). Data represent the mean±SEM of 3-6 mice each group. **E)** Representative 10x image of paraffine embedded KP-OVA tumor tissue labelled with Ly6G antibody (brown) and the corresponding 40 x magnification. Scale bars (100µm) are showed. **F)** Representative cryo-sections showing Ly6G (green) and SiglecF (red) in tumor nodules and the corresponding magnification. Scale bars (50 or 10µm) are indicated.

Immunohistochemistry readily detected a dense infiltrate of Ly6G^+^ cells located deeply into tumor nodules (Fig 1E), the majority of which stained also positive for SiglecF (Fig 1F).

### CXCL5 is highly expressed by mouse lung tumors and correlates to neutrophil infiltrate in human cancers

To explore the mechanism controlling accumulation of neu in lung tumor tissues we profiled normal and tumor lung tissues using a gene array of chemokines and chemokine receptors. KP-OVA lung tumor tissues displayed a 20-fold and 10-fold upregulation of *Cxcl5* and *Cxcl9* transcripts, respectively, and a slight induction of other chemokines implicated in monocytes and neu recruitment (Fig 2A). The CXCL5 chemokine binds to CXCR2 and it was previously associated to neu recruitment in inflammation and cancer (33-35). Fractionation of tumor tissues into CD45 positive and negative fractions showed that *Cxcl5* expression maps to cancer or stromal cells and not to infiltrating immune cells (Fig 2B). *Ex-vivo* analysis confirmed high *Cxcl5* production by KP cells and low expression in the other cell lines tested, including Lewis Lung carcinoma cells (3LL) (Fig 2C). We conclude that *Cxcl5* in KP tumor tissues derives primarily by a cancer-cell autonomous mechanism, possibly downstream mutated Kras (36).

**Figure 2.**
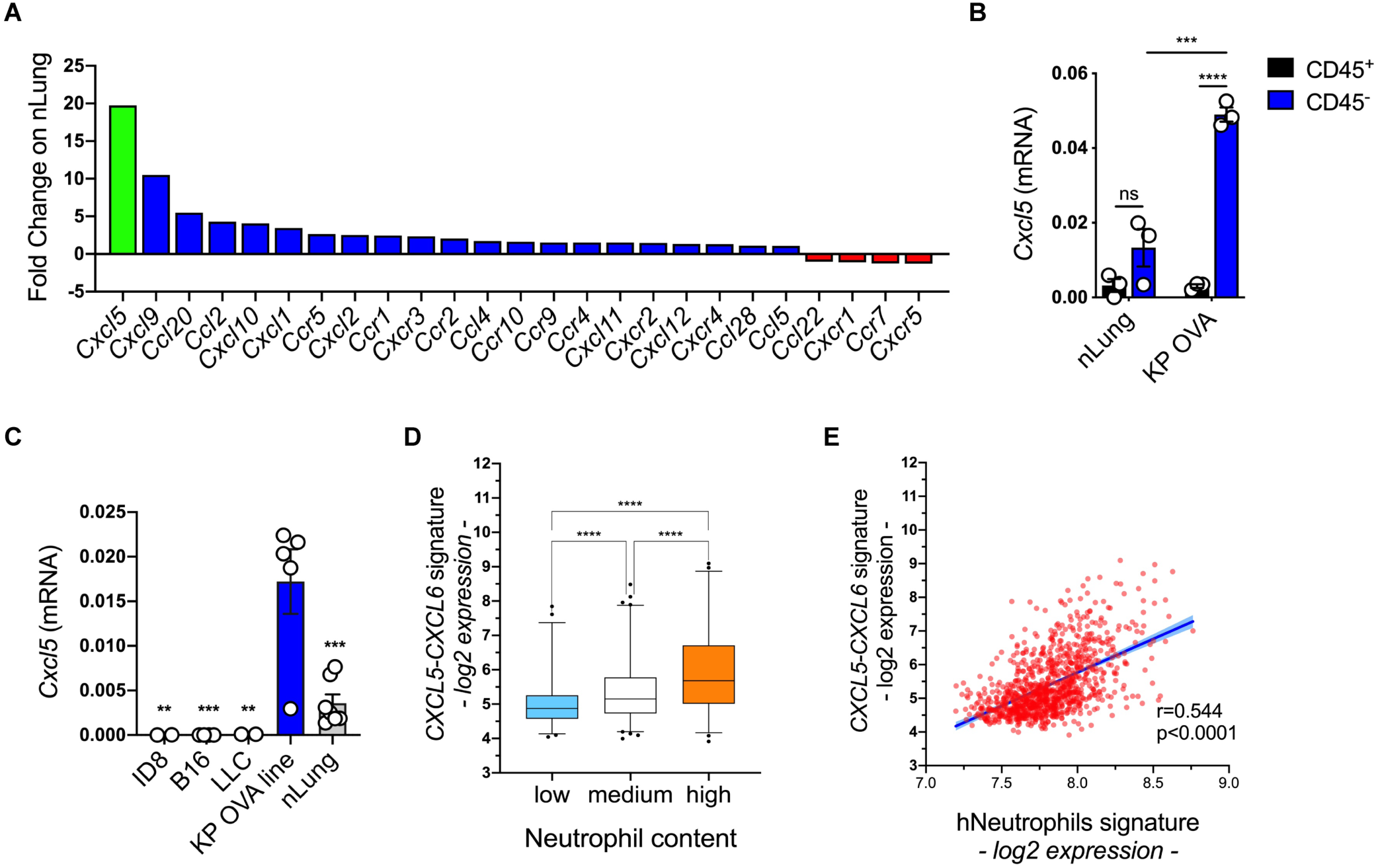
CXCL5 is highly expressed in mice tumors and human LUAD. **A)** Differential expression of chemokines and chemokine receptors in established KP-OVA lung tumor tissues over normal lung tissues. Data shows a representative of 2 independent experiments. **B)** Quantitative qRT-PCR of *Cxcl5* mRNA in CD45-positive or CD45-negative fractions isolated from nLung or KP OVA tumor bearing lungs. Graph shows mean±SEM of 3 replicates, significance was determined by one-way ANOVA with ***p≤0.001; ****p≤0.0001. **C)** Quantitative qRT-PCR of *Cxcl5* mRNA in different tumor cells and nLung was evaluated in 2-7 independent RNA extraction. Graph shows mean±SEM, significance over KP OVA cell line was determined by one-way ANOVA with **p≤0.01; ***p≤0.001. **D)** Patients from the compendium in Supplementary Table1, were stratified based on low, medium or high neu content based on CIBERSORT and correlated to the level of expression of a *CXCLs* chemokine (*CXCL5 and CXCL6*). **E)** Correlation between a human neu (hNeutrophils) gene signature and expression of *CXCLs* genes within the Compendium in Supplementary Table 1.

To validate these observations in the human context, we evaluated expression of *CXCL5*, and the highly homologous *CXCL6*, across the LUAD compendium. Patients were deconvolved using the CIBERSORT algorithm and divided into 3 categories, presenting low, medium and high neu density. Interestingly, we observed a high positive correlation between high chemokine expression and neu content (Fig 2D). Moreover, expression of tumor tissue neutrophils gene transcripts associated to bad prognosis (Fig 1A), strongly correlated to *CXCL5/6* expression in LUAD (Fig 2E). These data suggest that in humans, as in mice, enrichment of neu in lung cancer tissues is associated to overexpression of *CXCL5*.

### CXCL5 deletion inhibits accumulation of tumor associated SiglecF^high^ neu

We next targeted expression of CXCL5 in KP-OVA cells by CRISPR/CAS9 genome editing to assess its impact on accumulation of neu in lung tumors tissues. WT and KO^CXCL5^ KP-OVA clones were selected by ELISA and validated for equal OVA expression and *in vitro* growth rate (Fig 3A and S2A-C). Lung tumors formed by KP-OVA WT cells showed the expected increase in *Cxcl5* transcripts whereas lungs of mice injected with KP-OVA KO^CXCL5^ cells showed only a minor increase both in early and established tumors, confirming efficient deletion and indicating that the chemokine is produced primarily by cancer cells and not by stromal cells conditioned by tumor environmental factors (Fig 3B). Interestingly, KO^CXCL5^ tumors showed only a minor accumulation of neu both at early and advanced stages, with low SiglecF^high^ expression (Fig 3C). Immunohistochemistry confirmed lack of Ly6G^+^ cells in nodules of KO^CXCL5^ challenged mice, both at initial stages and in more advanced tumors (Fig 3D). Lack of CXCL5 secretion by KP-OVA cells had no impact on the frequency of circulating and splenic neu whereas it resulted in higher neu content within the bone marrow, likely reflecting reduced mobilization (S2D-F).

**Figure 3.**
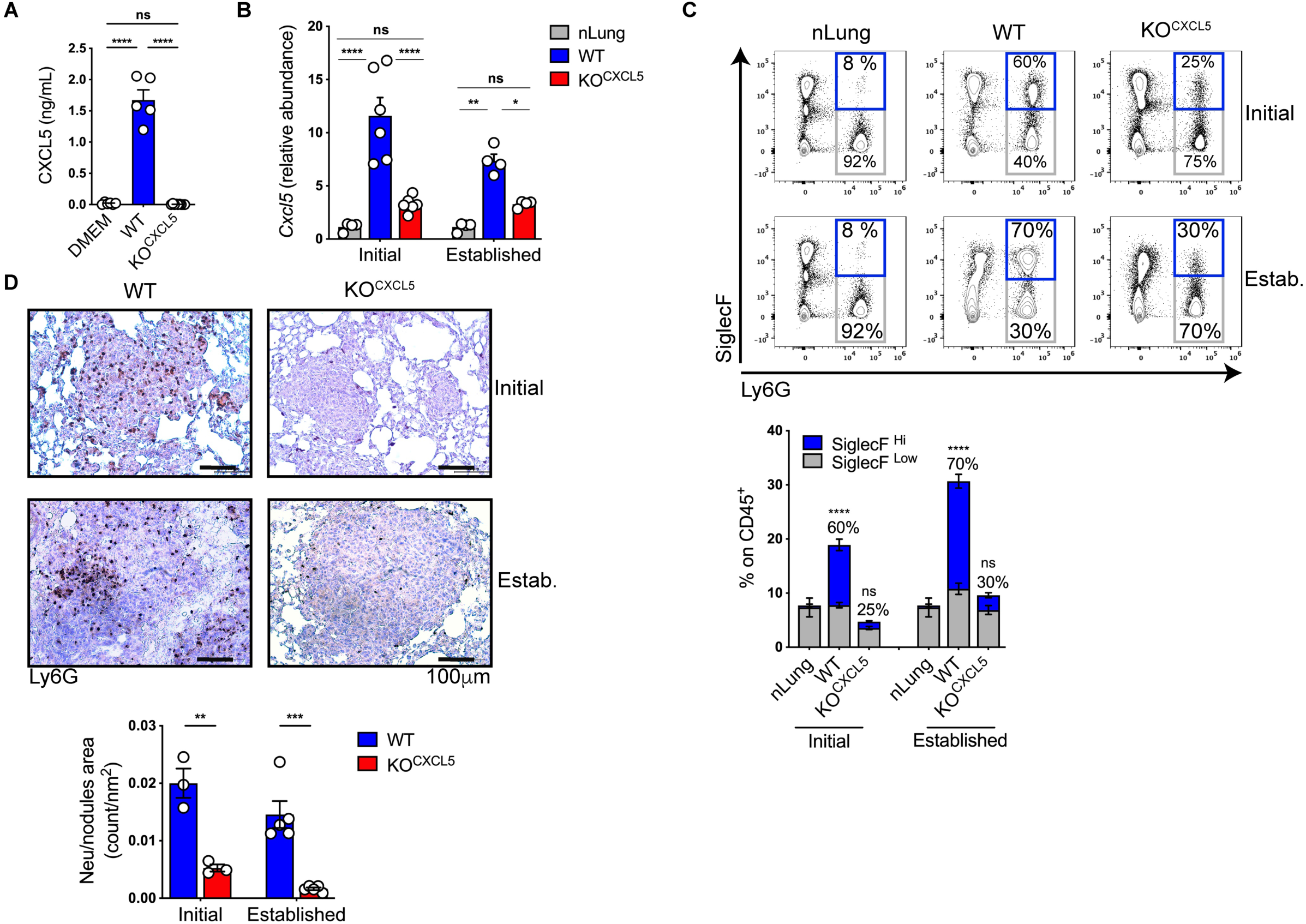
CXCL5 deficiency in immunogenic KP cells results in a selective and efficient inhibition of SiglecF^high^-neu recruitment in lung. **A)** The production of CXCL5 in the supernatant of KP-OVA WT and KO^CXCL5^ cells was measured by ELISA. Data represent the mean±SEM of 5-9 independent measurements. Significance was determined by a one-way ANOVA with ****p≤0.0001. **B-D)** Mice were challenged with PBS, KP-OVA WT and KO^CXCL5^. Lung tissues were harvested after 9 (initial) or 18 (established) days for analysis. **B)** Relative abundance of *Cxcl5* transcripts in total lung were evaluated by qRT-PCR. Data are expressed as fold induction over nLung. Data represent the mean±SEM of 4-6 independent RNA extraction. Significance was determined by two-way ANOVA with *p≤0.05, **p≤0.01, ****p≤0.0001. **C)** Representative flow cytometry dot plots of SiglecF^high/low^ expression on neu (gated on CD45^+^ cells). Number in quadrants refer to percentages of SiglecF^+^ on CD11b^+^/Ly6G^+^ cells. Bars on the right are mean of frequencies (%) of neu on live CD45^+^ lung cells. The fractions of SiglecF^High^ and SiglecF^Low^ within neu is indicated. Data represent the mean±SEM of 2 independent experiment, 3-4 mice each group. **D)** Representative IHC tissue sections of initial or established nodules labelled with Ly6G antibody (brown dots). Scale bars (100µm) are indicated. Graph on the right represents quantification of Ly6G^+^ on nodule area (count/nm^2^). Data are mean±SEM of 3-5 mice each group. Significance was determined by a two-way ANOVA with **p≤0.01; ***p≤0.001.

To ascertain that lack of neu recruitment in KO^CXCL5^ tumors is causally linked to chemokine expression, we reintroduced CXCL5 expression by lentiviral transduction, creating KO^CXCL5(lenti-^ _CXCL5)_ or control KO^CXCL5(lenti-vec)^ (Fig S3A). Cells were validated for CXCL5 expression *in vitro* and in lung tissues upon challenge (Fig S3B,C). Importantly, recruitment of neu was restored in the TME of KO^CXCL5(lenti-CXCL5)^ tumors, but not in control KO^CXCL5(lenti-vec)^, indicating that chemokine expression alone is sufficient to induce neu influx (Fig S3D). Notably, neu were highly recruited in KO^CXCL5(lenti-CXCL5)^ tumors but the fraction of SiglecF^high^ expressing cells was diminished as compared to WT. This finding suggests that CXCL5 overexpression accelerates recruitment of immature neu that are not yet fully converted into mature-aged SiglecF expressing neu, in line with recent reports pointing to local differentiation, aging and acquisition of SiglechF expression in lung tumor tissues (24, 25).

### Blocking accumulation of SiglecF^high^ neu improves activation of CD8 T cells

Multiple reports documented the T cell suppressive potential of circulating and splenic neutrophils isolated from tumor bearing hosts, in various cancer types, including lung cancer (9, 22, 37-39). The suppression of T cells in relevant, not ectopic, tumor tissues by well-defined specific neu subsets is less documented. Given that CXCL5 deletion in KP cells abrogates accumulation of SiglecF^high^ neu in lung cancer tissues we next sought to assess the consequences on tumor specific T cell responses, locally in lung tumor tissues. Animals were challenged with WT or KO^CXCL5^ KP-OVA cells and tissues were harvested at a time point corresponding to the peak of the T cell response in this model (30). Interestingly, the fraction of endogenous, OVA specific CD8 T cells in the lungs of animals challenged with KO^CXCL5^cancer cells was significantly higher (3-fold) than in WT tumors (Fig 4A). We used three combinations of markers to define the status of lung infiltrating T cells, as activated and functional (PD-1^+^LAG3^+^TIM3^+^), effector/memory (CD62L^-^CD44^+^), and tumor specific not exhausted (Tbet^+^EOMES^-^). All three parameters were statistically higher in CD8 T cells infiltrating KO^CXCL5^ lung tumors (Fig 4B). Importantly, ex-vivo restimulation of lung CD8 T cells with OVA class-I peptide to detect IFN-γ production, showed a 2-fold increase in the lungs of animals carrying KO^CXCL5^ tumors (Fig 4C), indicating enhanced cytotoxic functions. We observed a slight enhancement in the fraction of CD44^+^ CD8 T cells also in LN draining KO^CXCL5^ tumors, but no differences in markers of activation, effector memory or IFN-γ producing CD8 cells between the tumor genotypes (Fig S4A,B). We did not find any major difference in expression of activation markers in CD4 T cells infiltrating the lung or in tumor draining LNs (Fig S4C,D). These data are consistent with the notion that SiglecF^high^ neu suppress CD8 responses locally in the lung, whereas priming in LNs, where SiglecF^high^ neu are less abundant, is not primarily affected. To further establish whether SiglecF^high^ neu inhibit activation of tumor specific CD8 T cells or interfere with priming, we transferred OVA specific CD8 T cells (OTI) in tumor bearing mice. The extent of IFNγ production by OTI in the lung was higher in KO^CXCL5^ tumors, suggesting that SiglecF^high^ neu suppress ongoing responses (Fig 4D). To corroborate the impact of neu on local CD8 responses in the lung we performed as well depletion by classical Ly6G antibodies (3, 40). Depletion was efficient in the blood, however, flow cytometry and immunohistochemistry showed a large fraction of neu expressing SiglecF persisting in lung tumor tissues which precluded further analysis of T cell function (41)(Fig S5A-C). Collectively, these data show that SiglecF^high^ neu accumulating in lung tumor tissues inhibit locally anti-tumoral effector CD8 T cell.

**Figure 4.**
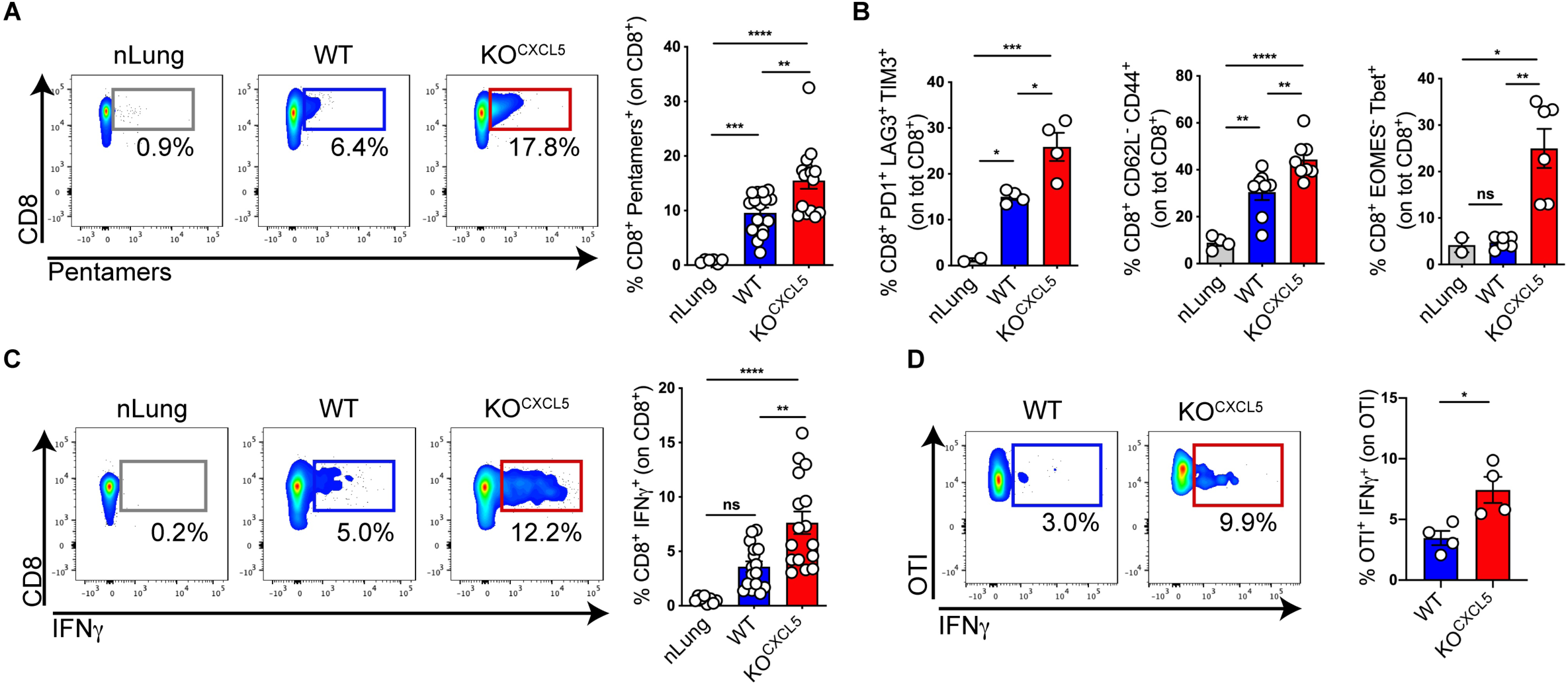
Lack of SiglecF^high^ neu promotes expansion and activation of CD8 T cells. **A-C)** Mice were challenged with WT or KO^CXCL5^ KP-OVA cells and endogenous T cell responses were analysed at early time point. **A)** Representative dot-plot showing OVA class-I pentamers labelling on lung CD8^+^ T cells. The fraction of CD8^+^pentamer^+^ over total CD8^+^ is plotted and quantified. Data are mean±SEM of 4 independent experiments with 4 mice/group. Significance was determined by one-way ANOVA with **p≤0.01, ***p≤0.001, ****p≤0.0001. **B)** Expression of T cell activation markers on CD8^+^ T cells in the lungs of control and tumor bearing animals. Frequencies of activated CD8^+^ PD1^+^TIM3^+^LAG3^+^ (left) memory CD8^+^CD62L^-^CD44^+^ (middle) and effector CD8^+^EOMES^-^Tbet^+^ (right) expressed as % on total CD8^+^ T cells are shown. Data represent mean±SEM of 2-8 mice/group. Significance was determined by one-way ANOVA with *p≤0.05, **p≤0.01, ***p≤0.001, ****p≤0.0001. **C)** Representative dot plot and quantification of IFNγ - producing lung CD8^+^ T cells. Data represent mean±SEM of 4 independent experiments with 2-4 mice each group. Significance was determined by one-way ANOVA with **p≤0.01, ****p≤0.0001. **D)** Mice received CFSE-labelled OVA-specific CD8^+^ T cells (OTI), 9 days after challenge with WT or KO^CXCL5^ KP OVA cells. OTI activation was measured by intracellular IFNγ staining. Representative dot plot and quantification of the fraction of IFNγ ^+^ OTI on total OTI, in tumor bearing lungs 2 days after transfer. Data represent mean±SEM of 4 mice each group. Significance was determined by unpaired t-test with *p≤0.05

### Neu density inversely correlates to CD8 T cell density in lung tumor tissues

Suppression of CD8 T cell by neu has been attributed to several pathways such as production of ROS, T cell antiproliferative molecules and nutrient deprivation (37). However, the relative spatial distribution of CD8 T cells and neu within tumor tissues remains poorly defined. Visualization of CD8 T cells within nodules of WT and KO^CXCL5^ tumors by immunohistochemistry showed a statistically significant increment in neu depleted nodules. To correlate the presence of CD8^+^ and neu we next performed co-labeling on cryosections. Unfortunately, triple labeling with SiglecF staining failed to unequivocally discriminate SiglecF^high^ neu from alveolar macrophages, so we focused on Ly6G to precisely identify and quantify neu. High magnification images of tumor nodules revealed several neu-CD8 synapses and some areas where CD8 T cells were surrounded by multiple neu, suggesting tight inter-cellular interactions. This is reminiscent of the recently described shielding of cancer cells by neutrophils extracellular traps, which may apply as well to sequestration of CD8 T cells (39) (Fig 5B). We further classified nodules as small (area<0,009mm^2^), medium (area between 0,0091mm^2^ and 0,02mm^2^) and large (area>0,021mm^2^) to quantify the number of CD8 T cells and neu within each nodule. Interestingly, CD8 T cells outnumbered neu in small nodules, were equal to neu in nodules of medium size and were underrepresented in larger nodules (Fig 5C), suggesting that inhibition may be mediated by physical interaction in tissues, rather than by systemic soluble signals.

**Figure 5.**
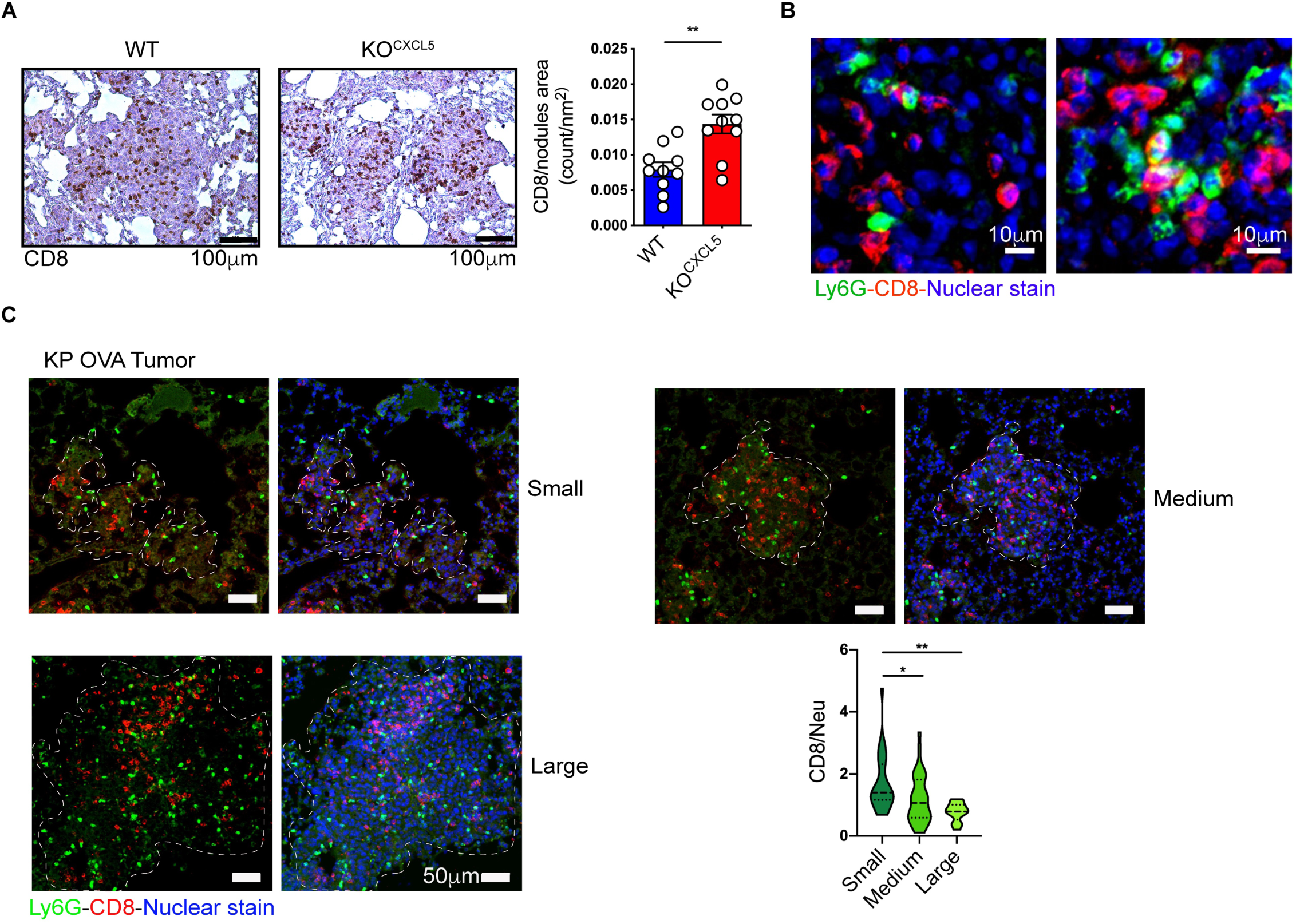
Spatial distribution of neu and CD8 T within tumor nodules. **A)** Representative images of WT or KO^CXCL5^ KP OVA lung tumor nodules labelled with anti CD8 antibodies (brown). Graph on the right shows quantification of the CD8^+^ infiltrate (CD8 count/nm^2^) in 2 independent experiments with 5 mice each group (mean±SEM). Significance was determined by an unpaired t-test with **p≤0.01. Scale bars (100µm) are indicated. **B)** High magnification examples of representative lung sections from WT KP OVA tumors co-labelled with antibodies to CD8 (red) and Ly6G (green) and counterstained with Hoechst to visualize nuclei. **C)** Nodules were classified into 3 categories based on the area (small<0,009 mm^2^, 0,0091 mm^2^ <medium<0,02 mm^2^, large>0,021 mm^2^). Scale bars (50µm) are indicated. On the right, quantification of the relative abundance of CD8 and neu within nodules of different sizes. Data are expressed as ratio of CD8/neu (mean±SEM of 90 nodules from 2 independent experiments with 2 mice each). Significance was determined by one-way ANOVA with *p≤0.05, **p≤0.01.

### SiglecF^high^ neu impair the efficacy of aPD-L1 checkpoint inhibitors

We next examined whether the observed enhancement of CD8 effector functions in KO^CXCL5^ tumors had an impact on tumor development. The size of tumor nodules and the number of proliferating tumor cells was evaluated in initial and established tumors. At early stages, nodules were significantly reduced in KO^CXCL5^ tumors and the fraction of proliferating cancer cells was diminished (Fig 6A). However, tumor growth was similar between the two genotypes at later time points, indicating that tumor containment by cytotoxic CD8 T cells was overcome at later stages. Expression of inhibitory ligands on tumor cells and tumor infiltrating myeloid cells is a key obstacle to spontaneous T cell responses to tumor antigens and neutrophils were previously documented to express PD-L1 in various settings (42, 43). Of note, we found that SiglecF^high^ neu express abundant PD-L1 molecules as compared to SiglecF^low^ neu, substantially contributing to increase the load of inhibitory ligands in the tumor microenvironment (Fig 6B,C). We thus reasoned that lowering PD-L1 burden by depleting SiglechF^high^ neu may favor effectiveness of checkpoint inhibitors. To test this hypothesis, we treated mice carrying WT or KO^CXCL5^ KP tumors with anti PD-L1 antibodies (Fig 6D). We established a common endpoint for all groups and the tumor burden was quantified as a proxy of treatment efficacy. PD-L1 blockade had no impact on the growth in WT KP tumors. In contrast, PD-L1 blockade effectively decreased the growth of KO^CXCL5^ tumors (Fig 6E-F). We concluded that accumulation of SiglechF^high^ neu with T cell suppressive activity restricts the efficacy of anti PD-L1 treatment in lung adenocarcinoma.

**Figure 6.**
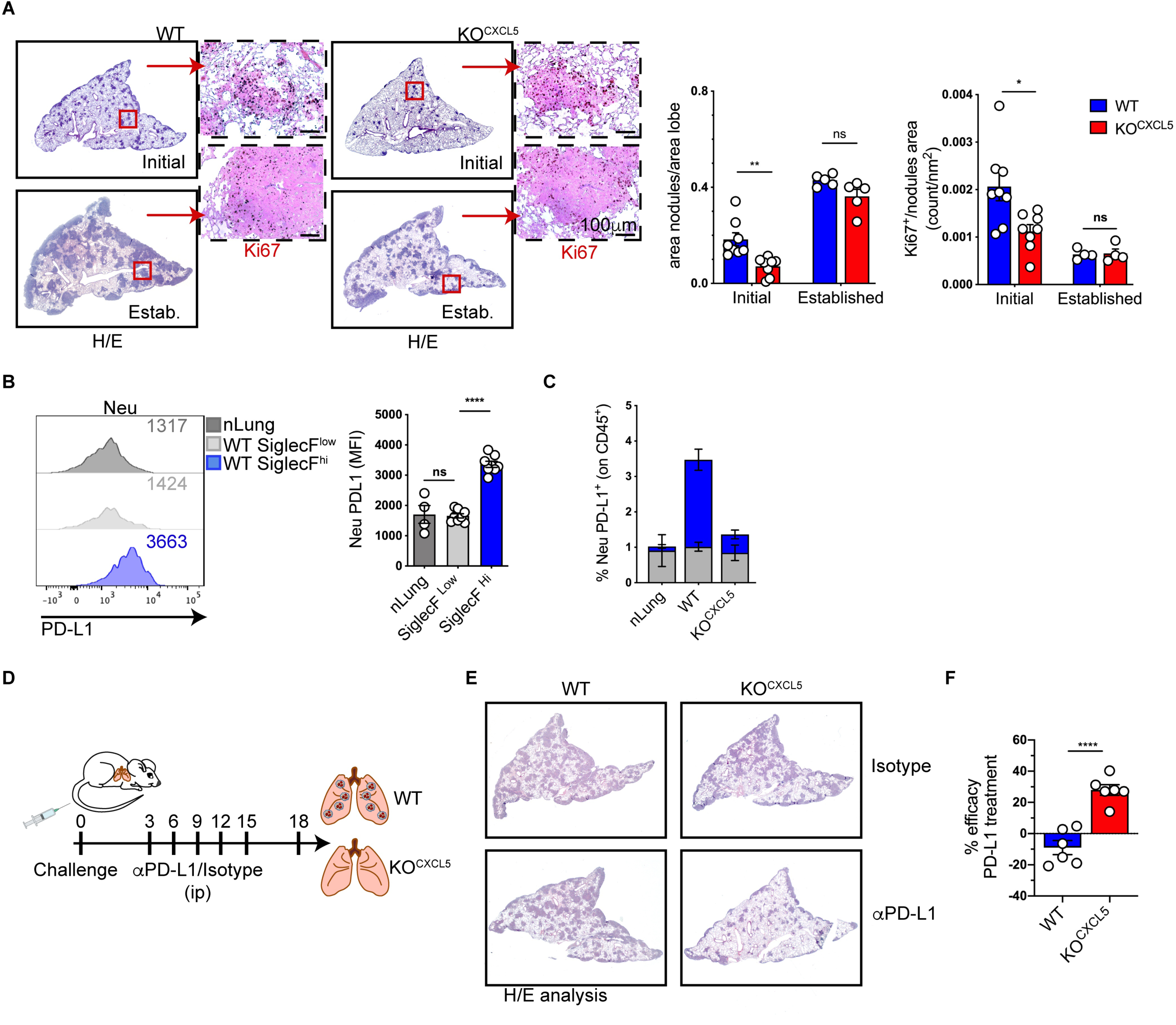
Targeting CXCL5-mediate neutrophils recruitment increase the effectiveness of αPD-L1 treatment. **A)** Mice were challenged with WT or KO^CXCL5^ KP-OVA cells and lungs were harvested after 9 or 18 days to score tumor burden (area nodules/area lobe) and cancer cell proliferation. Representative images of tumor nodules labelled by HE and Ki67 staining. Bars show quantification of tumor burden and fraction of Ki67+ cells/lobe area (nm^2^). Data are mean±SEM of 1 or 2 experiments with 4-5 mice each group. Significance was determined by two-way ANOVA with ns p>0.05, *p≤0.05, **p≤0.01. **B)** PD-L1 expression on lung infiltrating neu. Representative histogram of PD-L1 expression on neu from nLung or KP-OVA WT tumors. PD-L1 MFI (median fluorescence intensity) are indicated in the histogram and plotted on the right as mean±SEM of 2 experiments with 2-4 mice each group. Significance was determined by one-way ANOVA with ns p>0.05, ****p≤0.0001 **C)** Mice were challenged with WT or KO^CXCL5^ KP-OVA tumor cells and the frequency of PD-L1 expressing neu on total CD45^+^ cells was evaluated after 9 days. Data are mean±SEM of 2 independent experiments with 2-4 mice each group. **D)** Schematic representation of PD-L1 therapy in WT and KO^CXCL5^. Mice were treated with aPD-L1 or isotype control at day 3, 6, 9, 12 after tumor challenge. Lung tissue were harvested at day 18 to evaluate tumor burden by HE. **E)** Representative H&E staining of lung sections. **F)** Treatment efficacy is expressed as the percentage of reduction in tumor nodules over not treated controls. Data represent the mean±SEM of 2 experiments with 3 mice each group. Significance was determined by two-way ANOVA with ns p>0.05, *p≤0.05, ****p≤0.0001.

## Discussion

In this study we provide the first evidence that SiglecF^high^ neu accumulating in murine lung adenocarcinomas inhibit anti-cancer CD8 T cell responses and restrict the efficacy of PD-L1 blockade. These findings complement previous reports characterizing the origin and properties of SiglecF^high^ neu by uncovering their T cells suppressive potential. Importantly, our study identifies CXCL5 as sufficient to attract SiglecF^high^ neu in KP lung murine tumors and show its correlation with neu gene signatures in human LUAD. SiglecF^high^ neu were initially characterized as a protumorigenic subtype with high ROS and VEGF-A production, arising as a consequence of altered osteoblastic activity in the bone marrow of lung tumor hosts (23). Subsequent studies demonstrated that SiglecF^high^ neu are mature, aged neu with an enhanced glucose metabolism, that differentiate in tumor tissues (25). Neutralization of systemic G-CSF blunted recruitment of neu and decreased the proportion of SiglecF^high^ neu, however the pathway that regulate neu accumulation in KP lungs remained elusive. Infiltration of tumor tissues by neu has been associated to deregulated expression of CXCR2 ligands such as CXCL1, CXCL5 and CXCL6 in hepato-cellular carcinoma (HCC), breast cancer and squamous lung cancer (44-46). *In vivo* studies directly associated mutations in KRAS and LBK1 to overexpression of CXCL5 and CXCL8 and recruitment of neu (36). Our results align with these previous findings showing that several ligands of CXCR2 are overexpressed in neu infiltrated KP lung tumor tissues, with CXCL5 showing the highest levels. Genetic deletion of CXCL5 and the complementary overexpression experiment demonstrate the non-redundant role of the chemokine in promoting accumulation of SiglecF^high^ neu in KP tumors. Interestingly, CXCL5 overexpression rescued neu recruitment and increased the proportion of SiglecF^low^ cells (Fig S3D), suggesting that the chemokine induces accumulation of young neu precursors, rather than controlling local conversion into SiglecF expressing neu. Transplantable KP lines were validated in several previous studies addressing the composition of the immune infiltrate and testing combinatorial therapy (20, 28). SiglecF^high^ neu accumulate also in autochtonous KP tumors (25), yet it remains to be explored whether recruitment in slow progressing lesions may rely on the same or a different set of chemokines. Human data somehow reinforce the role of *CXCL5*, and its homolog *CXCL6*, in controlling neu density in human LUAD, a finding reported also in the context of HCC (47), colon cancer (35), melanoma (34) and lung inflammation (48-50), where it was proposed to be related to lung cancer progression through various mechanisms (45, 51, 52).

Genetic deletion of CXCL5 in the OVA expressing KP line offered the opportunity to analyze CD8 T cell responses in a tumor microenvironment depleted in SiglechF^high^ neu, avoiding the limitations of poorly specific and inefficient antibody depletion. The increment in numbers and the improvement of effector functions in cancer specific CD8 T cells in CXCL5 deficient KP tumors, uncovered the immune suppressive potential of SiglechF^high^ neu, a feature that was not investigated in previous reports. Although the mechanism of T cell suppression was not addressed in this study, imaging of tumor nodules identified areas of tight neu-CD8 T cells synapses and a gradual inversion of the CD8/neu ratio in small nodules as compared to larger nodules, suggesting replacement of CD8 by incoming neu. Expression of PD-L1 on neu was previously identified in HCC (42), gastric cancers (43) and on activated CD14^+^ neu infiltrating mouse ectopic tumors (22) and we here show that it is higher on SiglecF^high^ neu as compared to SiglecF^low^ neu. In lung cancer, high PD-L1 expression in the tumor and a high neu to lymphocyte ratio are two major predictors of response to immunotherapy (53-55). However, the contribution of PD-L1 expressed by neu to immune suppression and immunotherapy resistance has not formally addressed. Our results show that depleting SiglecF^high^ neu makes refractory KP tumor responsive to anti PD-L1 treatment. This may be explained by a quantitative effect by which SiglecF^high^ neu depletion lower the load of PD-L1 molecules to a level that allows effective blockade. Alternatively, other inhibitory signals arising from SiglecF^high^ neu and unrelated to PD-L1 expression, counteract the efficacy of checkpoint inhibition. Treatment of tumors with blockers of neu infiltration using CXCR2 inhibitors alone or in combination with checkpoint inhibitors are actively investigated in preclinical models and clinical trials. In conclusion our study uncovers that SiglechF^high^ neu accumulating in lung cancer tissues suppress spontaneous and immunotherapy induced anti-cancer T cells responses and suggest interference with the CXCL5axis as a way to improve the efficacy of checkpoint blockade.

## Acknowledgments

We thank Silvano Piazza and Alfonso Esposito, ICGEB Trieste for initial support on bioinformatic analysis and Luciano Morosi, ICGEB Trieste, for support on statistical analysis. We thank Prof. Annalisa del Prete, University of Brescia, for helpful discussions and advises on the KP mouse model. We thank Prof Mauro Giacca, ICGEB Trieste for providing the CXCL5 coding sequence.

## Contributors

FS, GMP and TB performed experiments. FS and GMP analysed data, prepared figures and contributed to manuscript writing. AC provided CXCL5 expression construct. SB analyzed human data. NC set up the tumor model and generated initial observations. FB ideated and supervised the study and wrote the manuscript.

## Funding

This work was supported by an AIRC grant to FB (IG 21636). FS was supported by an ICGEB Arturo Falaschi pre-doctoral fellowships.

## Competing interests

The authors have no conflict of interest.

## Ethical approval

The study was approved by International Centre for Genetic Engineering and Biotechnology (ICGEB) board for animal welfare and authorized by the Italian Ministry of Health (approval number 1133/2020-PR, issued on 12/11/2020). Animal care and treatment were conducted with national and international laws and policies (European Economic Community Council Directive 86/609; OJL 358; December 12, 1987). All experiments were performed in accordance with the Federation of European Laboratory Animal Science Association (FELASA) guidelines

## Data availability statement

11 data relevant to the study are included in the article or uploaded as supplementary information. The datasets used and analyzed during the current study are included in the article or uploaded as supplementary Information. They are available from the corresponding author on reasonable request.

---

Genes used to determine the hNeutrophils signature in Figure 1A and 2E.

| hNeutrophils |  |  |  |
| --- | --- | --- | --- |
| PI3 | DGAT2 | MXD1 | XPO6 |
| CXCL8 | LCP2 | MMP25 | DUSP1 |
| G0S2 | FPR2 | IER3 | TNFRSF1A |
| CXCR2 | GCA | LIMK2 | MBOAT7 |
| S100A8 | PLEK | PPIF | KCNJ15 |
| FCGR3B | CXCL1 | SMIM25 | ACSL1 |
| IL1R2 | AQP9 | CYSTM1 | TDP2 |
| PROK2 | AL356273.3 | FOS | RIPOR2 |
| IFITM2 | USP10 | LINC01001 | RALGDS |
| CSF3R | SLPI | MME | ATP2B1-AS1 |
| TREM1 | CCRL2 | SAT1 | UBE2B |
| AL034397.3 | SOD2 | ADGRG3 | CPD |
| AC245128.3 | FFAR2 | MTHFS | BASP1 |
| OSM | HCAR3 | AC023157.3 | PADI4 |
| IVNS1ABP | VNN2 | CEBPB | DOK3 |
| IL1RN | HCAR2 | CDA | MAP1LC3B2 |
| BCL2A1 | FPR1 | CCL4L2 | C15orf48 |
| S100P | CMTM2 | CYP4F3 | H3F3A |
| S100A9 | NAMPT | MAP1LC3B | ETS2 |
| S100A12 | CXCR1 | LITAF | IFIT3 |
| KATNBL1 | LRG1 | IL1A | NSMAF |
| ALOX5AP | TNFAIP6 | RILPL2 | LRRK2 |
| TRIB1 | PHACTR1 | LINC01506 | RNF24 |
| CLEC4E | ADM | CLEC4A | MSRB1 |
| ANP32A | FGR |  |  |

## Notes

### Competing Interest Statement

The authors have declared no competing interest.

## References

1. Houghton AM, Rzymkiewicz DM, Ji H, Gregory AD, Egea EE, Metz HE, et al. Neutrophil elastase-mediated degradation of IRS-1 accelerates lung tumor growth. Nat Med. 2010;16(2):219–23.

2. Butin-Israeli V, Bui TM, Wiesolek HL, Mascarenhas L, Lee JJ, Mehl LC, et al. Neutrophil-induced genomic instability impedes resolution of inflammation and wound healing. J Clin Invest. 2019;129(2):712–26.

3. Coffelt SB, Kersten K, Doornebal CW, Weiden J, Vrijland K, Hau CS, et al. IL-17-producing gammadelta T cells and neutrophils conspire to promote breast cancer metastasis. Nature. 2015;522(7556):345–8.

4. Wculek SK, Malanchi I. Neutrophils support lung colonization of metastasis-initiating breast cancer cells. Nature. 2015;528(7582):413–7.

5. Albrengues J, Shields MA, Ng D, Park CG, Ambrico A, Poindexter ME, et al. Neutrophil extracellular traps produced during inflammation awaken dormant cancer cells in mice. Science. 2018;361(6409).

6. Nozawa H, Chiu C, Hanahan D. Infiltrating neutrophils mediate the initial angiogenic switch in a mouse model of multistage carcinogenesis. Proc Natl Acad Sci U S A. 2006;103(33):12493–8.

7. Albini A, Bruno A, Noonan DM, Mortara L. Contribution to Tumor Angiogenesis From Innate Immune Cells Within the Tumor Microenvironment: Implications for Immunotherapy. Front Immunol. 2018;9:527.

8. Rice CM, Davies LC, Subleski JJ, Maio N, Gonzalez-Cotto M, Andrews C, et al. Tumour-elicited neutrophils engage mitochondrial metabolism to circumvent nutrient limitations and maintain immune suppression. Nat Commun. 2018;9(1):5099.

9. Veglia F, Tyurin VA, Blasi M, De Leo A, Kossenkov AV, Donthireddy L, et al. Fatty acid transport protein 2 reprograms neutrophils in cancer. Nature. 2019;569(7754):73–8.

10. Glodde N, Bald T, van den Boorn-Konijnenberg D, Nakamura K, O’Donnell JS, Szczepanski S, et al. Reactive Neutrophil Responses Dependent on the Receptor Tyrosine Kinase c-MET Limit Cancer Immunotherapy. Immunity. 2017;47(4):789–802 e9.

11. Ponzetta A, Carriero R, Carnevale S, Barbagallo M, Molgora M, Perucchini C, et al. Neutrophils Driving Unconventional T Cells Mediate Resistance against Murine Sarcomas and Selected Human Tumors. Cell. 2019;178(2):346–60 e24.

12. Singhal S, Bhojnagarwala PS, O’Brien S, Moon EK, Garfall AL, Rao AS, et al. Origin and Role of a Subset of Tumor-Associated Neutrophils with Antigen-Presenting Cell Features in Early-Stage Human Lung Cancer. Cancer Cell. 2016;30(1):120–35.

13. Templeton AJ, McNamara MG, Seruga B, Vera-Badillo FE, Aneja P, Ocana A, et al. Prognostic role of neutrophil-to-lymphocyte ratio in solid tumors: a systematic review and meta-analysis. J Natl Cancer Inst. 2014;106(6):dju124.

14. Gentles AJ, Newman AM, Liu CL, Bratman SV, Feng W, Kim D, et al. The prognostic landscape of genes and infiltrating immune cells across human cancers. Nat Med. 2015;21(8):938–45.

15. Kargl J, Busch SE, Yang GH, Kim KH, Hanke ML, Metz HE, et al. Neutrophils dominate the immune cell composition in non-small cell lung cancer. Nat Commun. 2017;8:14381.

16. Rapoport BL, Theron AJ, Vorobiof DA, Langenhoven L, Hall JM, Van Eeden RI, et al. Prognostic significance of the neutrophil/lymphocyte ratio in patients undergoing treatment with nivolumab for recurrent non-small-cell lung cancer. Lung Cancer Manag. 2020;9(3):LMT37.

17. Russo A, Russano M, Franchina T, Migliorino MR, Aprile G, Mansueto G, et al. Neutrophil-to-Lymphocyte Ratio (NLR), Platelet-to-Lymphocyte Ratio (PLR), and Outcomes with Nivolumab in Pretreated Non-Small Cell Lung Cancer (NSCLC): A Large Retrospective Multicenter Study. Adv Ther. 2020;37(3):1145–55.

18. Jiang T, Bai Y, Zhou F, Li W, Gao G, Su C, et al. Clinical value of neutrophil-to-lymphocyte ratio in patients with non-small-cell lung cancer treated with PD-1/PD-L1 inhibitors. Lung Cancer. 2019;130:76–83.

19. Mitchell KG, Diao L, Karpinets T, Negrao MV, Tran HT, Parra ER, et al. Neutrophil expansion defines an immunoinhibitory peripheral and intratumoral inflammatory milieu in resected non-small cell lung cancer: a descriptive analysis of a prospectively immunoprofiled cohort. J Immunother Cancer. 2020;8(1).

20. Zilionis R, Engblom C, Pfirschke C, Savova V, Zemmour D, Saatcioglu HD, et al. Single-Cell Transcriptomics of Human and Mouse Lung Cancers Reveals Conserved Myeloid Populations across Individuals and Species. Immunity. 2019;50(5):1317–34 e10.

21. Shaul ME, Eyal O, Guglietta S, Aloni P, Zlotnik A, Forkosh E, et al. Circulating neutrophil subsets in advanced lung cancer patients exhibit unique immune signature and relate to prognosis. FASEB J. 2020;34(3):4204–18.

22. Veglia F, Hashimoto A, Dweep H, Sanseviero E, De Leo A, Tcyganov E, et al. Analysis of classical neutrophils and polymorphonuclear myeloid-derived suppressor cells in cancer patients and tumor-bearing mice. J Exp Med. 2021;218(4).

23. Engblom C, Pfirschke C, Zilionis R, Da Silva Martins J, Bos SA, Courties G, et al. Osteoblasts remotely supply lung tumors with cancer-promoting SiglecF(high) neutrophils. Science. 2017;358(6367).

24. Pfirschke C, Engblom C, Gungabeesoon J, Lin Y, Rickelt S, Zilionis R, et al. Tumor-Promoting Ly-6G(+) SiglecF(high) Cells Are Mature and Long-Lived Neutrophils. Cell Rep. 2020;32(12):108164.

25. Ancey PB, Contat C, Boivin G, Sabatino S, Pascual J, Zangger N, et al. GLUT1 Expression in Tumor-Associated Neutrophils Promotes Lung Cancer Growth and Resistance to Radiotherapy. Cancer Res. 2021;81(9):2345–57.

26. Vafadarnejad E, Rizzo G, Krampert L, Arampatzi P, Arias-Loza AP, Nazzal Y, et al. Dynamics of Cardiac Neutrophil Diversity in Murine Myocardial Infarction. Circ Res. 2020;127(9):e232–e49.

27. Matsui M, Nagakubo D, Satooka H, Hirata T. A novel Siglec-F(+) neutrophil subset in the mouse nasal mucosa exhibits an activated phenotype and is increased in an allergic rhinitis model. Biochem Biophys Res Commun. 2020;526(3):599–606.

28. Maier B, Leader AM, Chen ST, Tung N, Chang C, LeBerichel J, et al. A conserved dendritic-cell regulatory program limits antitumour immunity. Nature. 2020;580(7802):257–62.

29. Dimitrova N, Gocheva V, Bhutkar A, Resnick R, Jong RM, Miller KM, et al. Stromal Expression of miR-143/145 Promotes Neoangiogenesis in Lung Cancer Development. Cancer Discov. 2016;6(2):188–201.

30. Caronni N, Piperno GM, Simoncello F, Romano O, Vodret S, Yanagihashi Y, et al. TIM4 expression by dendritic cells mediates uptake of tumor-associated antigens and anti-tumor responses. Nat Commun. 2021;12(1):2237.

31. Newman AM, Liu CL, Green MR, Gentles AJ, Feng W, Xu Y, et al. Robust enumeration of cell subsets from tissue expression profiles. Nat Methods. 2015;12(5):453–7.

32. DuPage M, Cheung AF, Mazumdar C, Winslow MM, Bronson R, Schmidt LM, et al. Endogenous T cell responses to antigens expressed in lung adenocarcinomas delay malignant tumor progression. Cancer Cell. 2011;19(1):72–85.

33. Wang L, Shi L, Gu J, Zhan C, Xi J, Ding J, et al. CXCL5 regulation of proliferation and migration in human non-small cell lung cancer cells. J Physiol Biochem. 2018;74(2):313–24.

34. Forsthuber A, Lipp K, Andersen L, Ebersberger S, Graña C, Ellmeier W, et al. CXCL5 as Regulator of Neutrophil Function in Cutaneous Melanoma. J Invest Dermatol. 2019;139(1):186–94.

35. Lin Y, Cheng L, Liu Y, Wang Y, Wang Q, Wang HL, et al. Intestinal epithelium-derived BATF3 promotes colitis-associated colon cancer through facilitating CXCL5-mediated neutrophils recruitment. Mucosal Immunol. 2021;14(1):187–98.

36. Wislez M, Fujimoto N, Izzo JG, Hanna AE, Cody DD, Langley RR, et al. High expression of ligands for chemokine receptor CXCR2 in alveolar epithelial neoplasia induced by oncogenic kras. Cancer Res. 2006;66(8):4198–207.

37. Condamine T, Dominguez GA, Youn JI, Kossenkov AV, Mony S, Alicea-Torres K, et al. Lectin-type oxidized LDL receptor-1 distinguishes population of human polymorphonuclear myeloid-derived suppressor cells in cancer patients. Sci Immunol. 2016;1(2).

38. Cheng Y, Li H, Deng Y, Tai Y, Zeng K, Zhang Y, et al. Cancer-associated fibroblasts induce PDL1+ neutrophils through the IL6-STAT3 pathway that foster immune suppression in hepatocellular carcinoma. Cell Death Dis. 2018;9(4):422.

39. Teijeira A, Garasa S, Gato M, Alfaro C, Migueliz I, Cirella A, et al. CXCR1 and CXCR2 Chemokine Receptor Agonists Produced by Tumors Induce Neutrophil Extracellular Traps that Interfere with Immune Cytotoxicity. Immunity. 2020;52(5):856–71 e8.

40. Zhu J, Powis de Tenbossche CG, Cane S, Colau D, van Baren N, Lurquin C, et al. Resistance to cancer immunotherapy mediated by apoptosis of tumor-infiltrating lymphocytes. Nat Commun. 2017;8(1):1404.

41. Boivin G, Faget J, Ancey PB, Gkasti A, Mussard J, Engblom C, et al. Durable and controlled depletion of neutrophils in mice. Nat Commun. 2020;11(1):2762.

42. He G, Zhang H, Zhou J, Wang B, Chen Y, Kong Y, et al. Peritumoural neutrophils negatively regulate adaptive immunity via the PD-L1/PD-1 signalling pathway in hepatocellular carcinoma. J Exp Clin Cancer Res. 2015;34:141.

43. Wang TT, Zhao YL, Peng LS, Chen N, Chen W, Lv YP, et al. Tumour-activated neutrophils in gastric cancer foster immune suppression and disease progression through GM-CSF-PD-L1 pathway. Gut. 2017;66(11):1900–11.

44. Acharyya S, Oskarsson T, Vanharanta S, Malladi S, Kim J, Morris PG, et al. A CXCL1 paracrine network links cancer chemoresistance and metastasis. Cell. 2012;150(1):165–78.

45. Mollaoglu G, Jones A, Wait SJ, Mukhopadhyay A, Jeong S, Arya R, et al. The Lineage-Defining Transcription Factors SOX2 and NKX2-1 Determine Lung Cancer Cell Fate and Shape the Tumor Immune Microenvironment. Immunity. 2018;49(4):764-79.e9.

46. Koyama S, Akbay EA, Li YY, Aref AR, Skoulidis F, Herter-Sprie GS, et al. STK11/LKB1 Deficiency Promotes Neutrophil Recruitment and Proinflammatory Cytokine Production to Suppress T-cell Activity in the Lung Tumor Microenvironment. Cancer Res. 2016;76(5):999–1008.

47. Zhou SL, Dai Z, Zhou ZJ, Wang XY, Yang GH, Wang Z, et al. Overexpression of CXCL5 mediates neutrophil infiltration and indicates poor prognosis for hepatocellular carcinoma. Hepatology. 2012;56(6):2242–54.

48. Nouailles G, Dorhoi A, Koch M, Zerrahn J, Weiner J, 3rd, Fae KC, et al. CXCL5-secreting pulmonary epithelial cells drive destructive neutrophilic inflammation in tuberculosis. J Clin Invest. 2014;124(3):1268–82.

49. Jovic S, Linge HM, Shikhagaie MM, Olin AI, Lannefors L, Erjefalt JS, et al. The neutrophil-recruiting chemokine GCP-2/CXCL6 is expressed in cystic fibrosis airways and retains its functional properties after binding to extracellular DNA. Mucosal Immunol. 2016;9(1):112–23.

50. Cheng Y, Ma XL, Wei YQ, Wei XW. Potential roles and targeted therapy of the CXCLs/CXCR2 axis in cancer and inflammatory diseases. Biochim Biophys Acta Rev Cancer. 2019;1871(2):289–312.

51. Eide HA, Knudtsen IS, Sandhu V, Løndalen AM, Halvorsen AR, Abravan A, et al. Serum cytokine profiles and metabolic tumor burden in patients with non-small cell lung cancer undergoing palliative thoracic radiation therapy. Adv Radiat Oncol. 2018;3(2):130–8.

52. Zhu YM, Bagstaff SM, Woll PJ. Production and upregulation of granulocyte chemotactic protein-2/CXCL6 by IL-1beta and hypoxia in small cell lung cancer. Br J Cancer. 2006;94(12):1936–41.

53. Gandhi L, Rodriguez-Abreu D, Gadgeel S, Esteban E, Felip E, De Angelis F, et al. Pembrolizumab plus Chemotherapy in Metastatic Non-Small-Cell Lung Cancer. N Engl J Med. 2018;378(22):2078–92.

54. Soyano AE, Dholaria B, Marin-Acevedo JA, Diehl N, Hodge D, Luo Y, et al. Peripheral blood biomarkers correlate with outcomes in advanced non-small cell lung Cancer patients treated with anti-PD-1 antibodies. J Immunother Cancer. 2018;6(1):129.

55. Ren F, Zhao T, Liu B, Pan L. Neutrophil-lymphocyte ratio (NLR) predicted prognosis for advanced non-small-cell lung cancer (NSCLC) patients who received immune checkpoint blockade (ICB). Onco Targets Ther. 2019;12:4235–44.

## References

Bild AH, Yao G, Chang JT, Wang Q et al. Oncogenic pathway signatures in human cancers as a guide to targeted therapies. Nature 2006 Jan 19;439(7074):353–7

Chitale D, Gong Y, Taylor BS, Broderick S, Brennan C, Somwar R, Golas B, Wang L, Motoi N, Szoke J, Reinersman JM, Major J, Sander C, Seshan VE, Zakowski MF, Rusch V, Pao W, Gerald W, Ladanyi M.. An integrated genomic analysis of lung cancer reveals loss of DUSP4 in EGFR-mutant tumors. Oncogene. 2009 Aug 6;28(31):2773–83

Hou J, Aerts J, den Hamer B, van Ijcken W et al. Gene expression-based classification of non-small cell lung carcinomas and survival prediction. PLoS One 2010 Apr 22;5(4):e10312

Kuner R, Muley T, Meister M, Ruschhaupt M et al. Global gene expression analysis reveals specific patterns of cell junctions in non-small cell lung cancer subtypes. Lung Cancer 2009 Jan;63(1):32–8

Nguyen DX, Chiang AC, Zhang XH, Kim JY, Kris MG, Ladanyi M, Gerald WL, Massagué J. WNT/TCF signaling through LEF1 and HOXB9 mediates lung adenocarcinoma metastasis. Cell. 2009 Jul 10;138(1):51–62

Okayama H, Kohno T, Ishii Y, Shimada Y et al. Identification of genes upregulated in ALK-positive and EGFR/KRAS/ALK-negative lung adenocarcinomas. Cancer Res 2012 Jan 1;72(1):100–11

Shedden K, Taylor JM, Enkemann SA et al. Gene expression-based survival prediction in lung adenocarcinoma: a multi-site, blinded validation study. Nat Med 2008 Aug;14(8):822–7

Yamauchi M, Yamaguchi R, Nakata A, Kohno T et al. Epidermal growth factor receptor tyrosine kinase defines critical prognostic genes of stage I lung adenocarcinoma. PLoS One 2012;7(9):e43923

Zhu CQ, Ding K, Strumpf D, Weir BA et al. Prognostic and predictive gene signature for adjuvant chemotherapy in resected non-small-cell lung cancer. J Clin Oncol 2010 Oct 10;28(29):4417–24

