## Supplementary materials for "SiglecF^high^ neutrophils in lung tumor tissues suppress local CD8 T cell responses and limit the efficacy of anti PD-L1 antibodies"

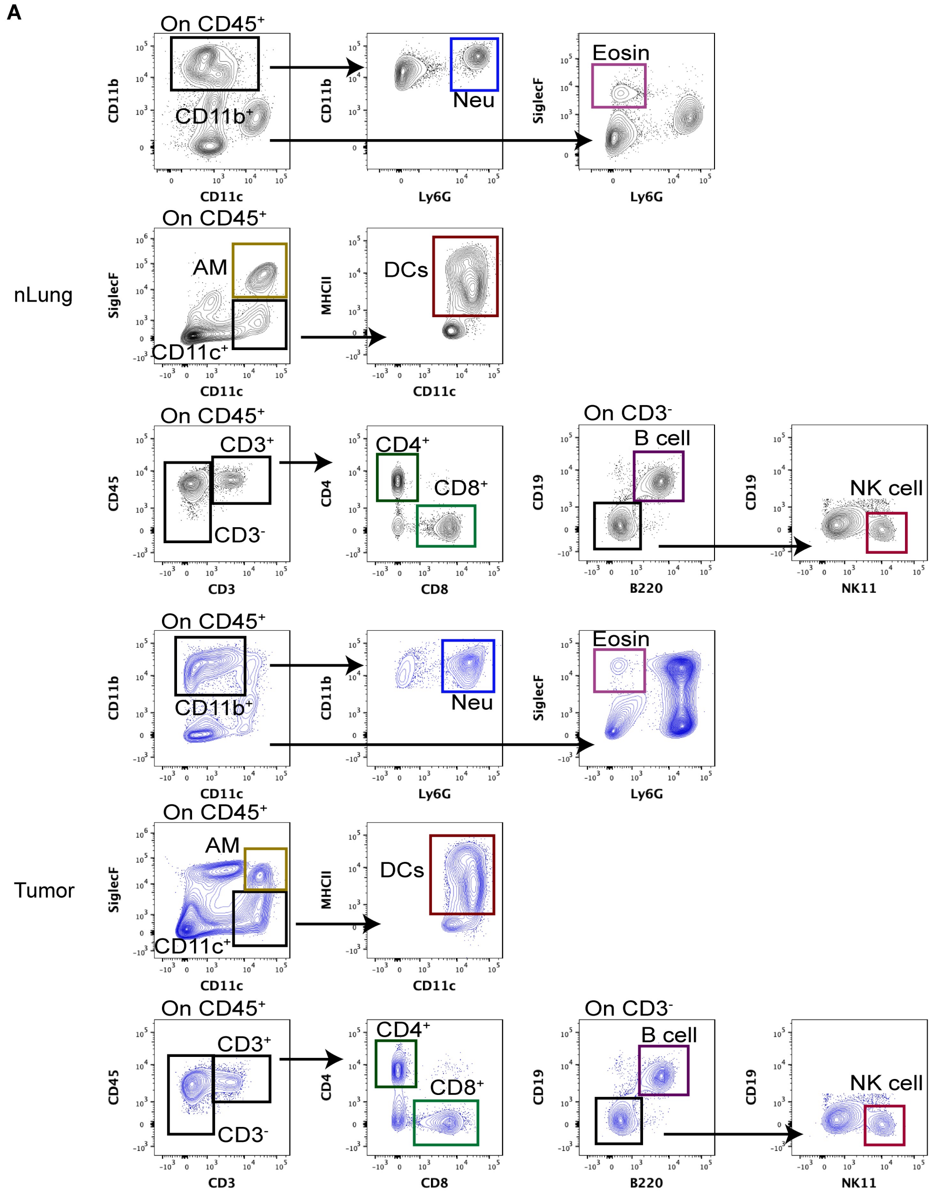


**Supplementary Fig. S1.**

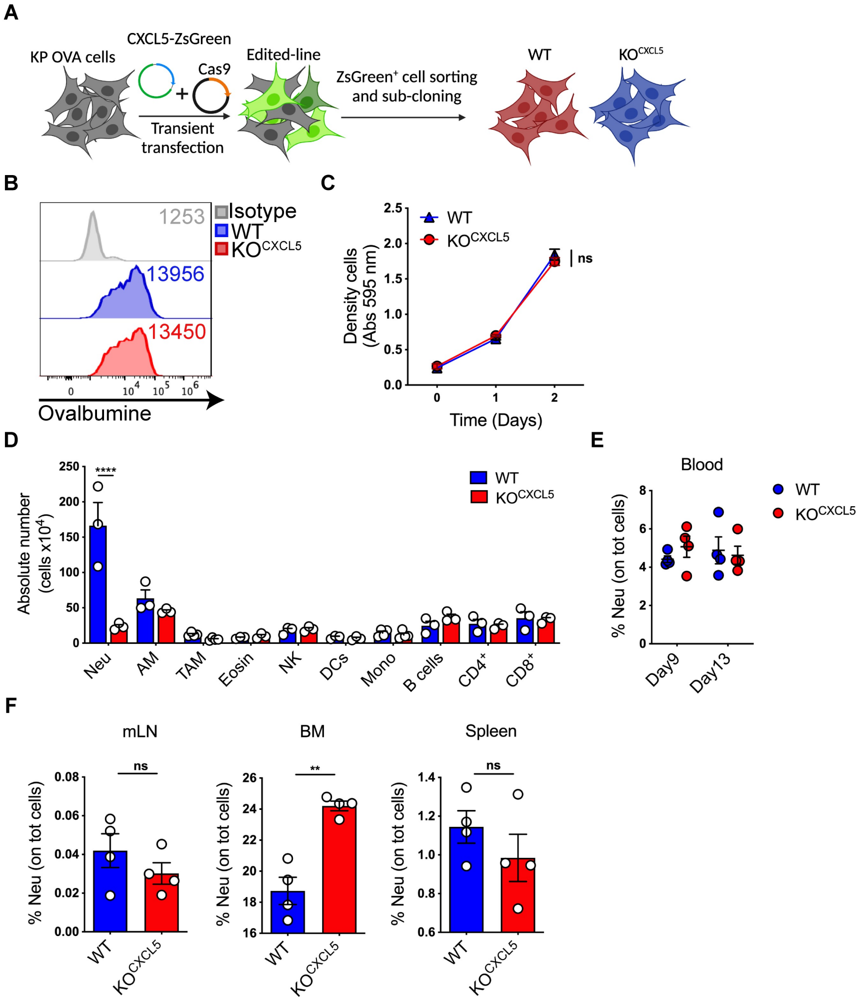


**Supplementary Fig. S2.**

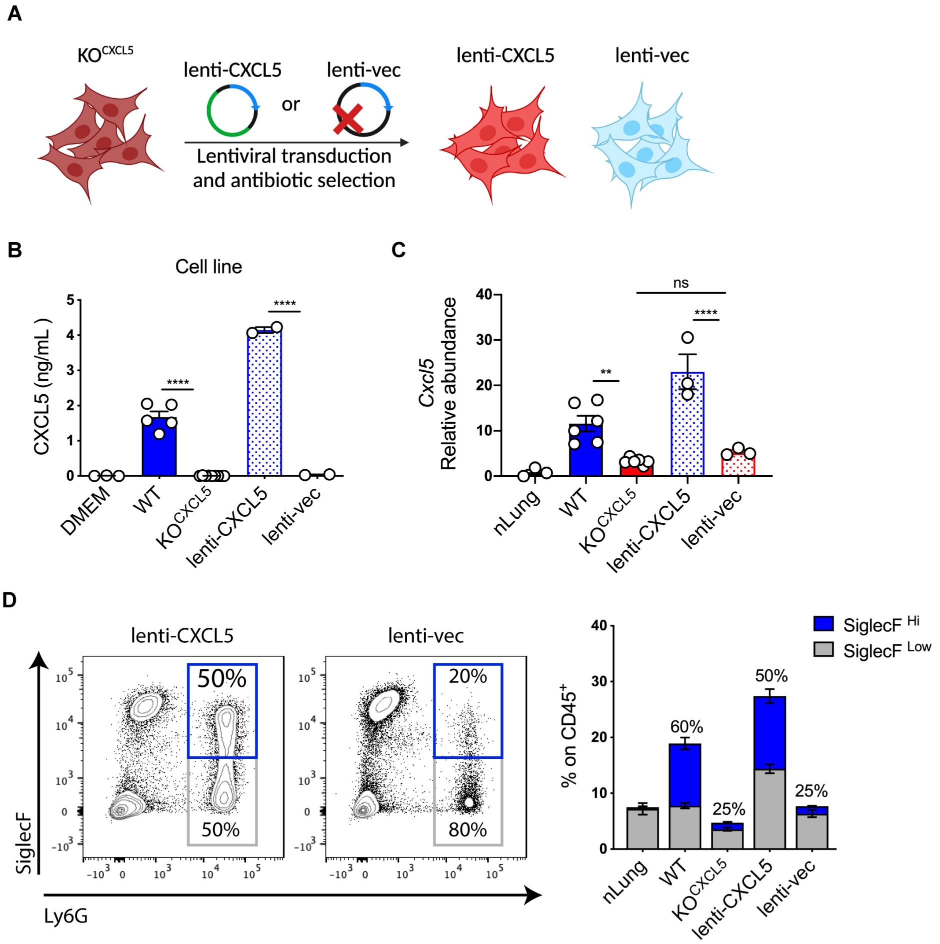


**Supplementary Fig. S3.**

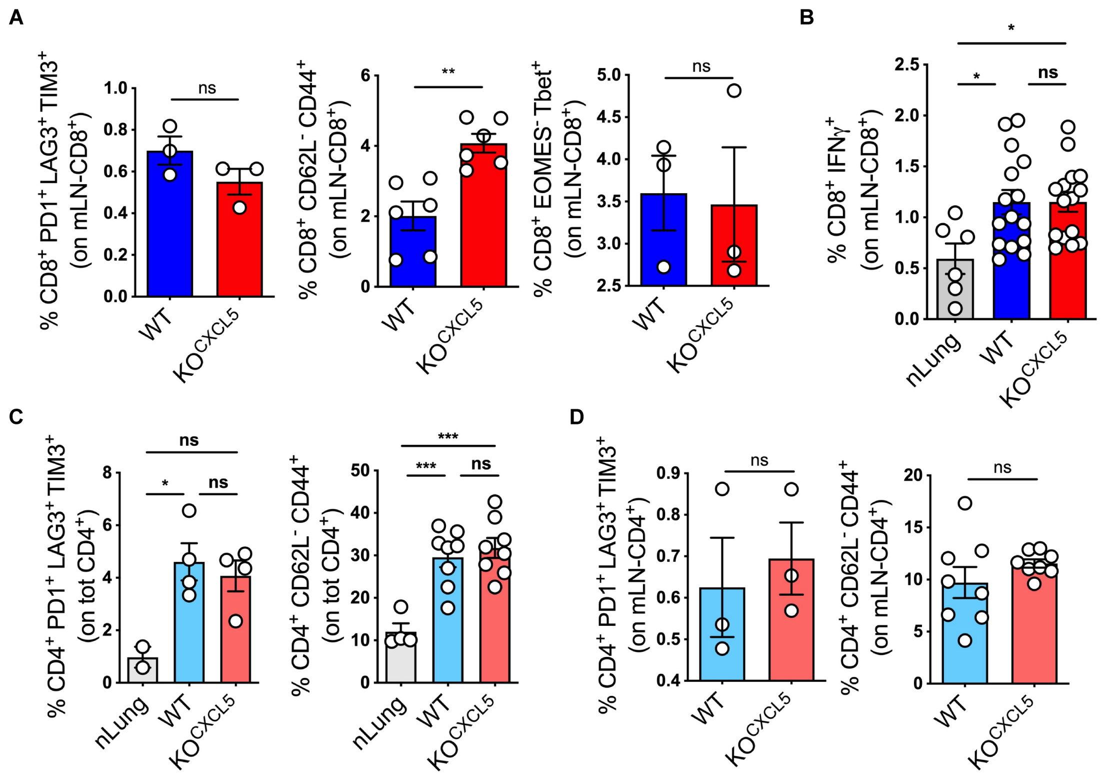


**Supplementary Fig. S4.**

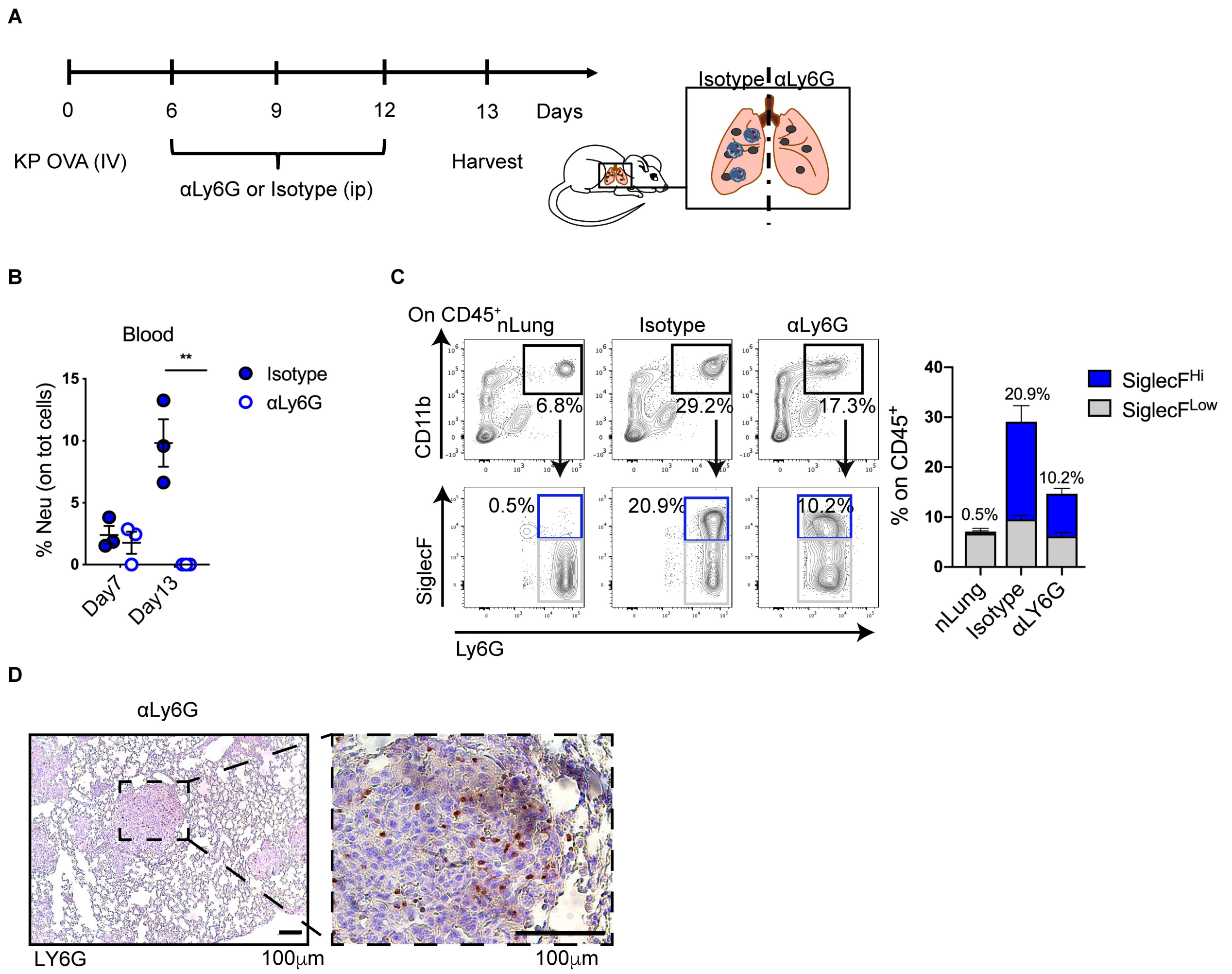


**Supplementary Fig. S5.**

### Supplementary Figure legends

**Supplementary Fig. S1. Immune subset identification in lung.** Gating strategy used for the identification of immune subsets in nLung (Upper) and tumor bearing lungs (lower). Flow cytometry plots showing the gating strategy to identify neu (=neutrophils), Eosin (=eosinophils), AM (=alveolar macrophages), DCs (=dendritic cells), NK (=natural killer cells), B cell, $CD4^+$, $CD8^+$.

**Supplementary Fig. S2. Generation and validation of $KO^{CXCL5}$ cell line. A)** Schematic representation of KP-OVA $KO^{CXCL5}$ cells generation. KP-OVA cells were transiently transfected with Cas9 and sgCXCL5-$ZsGreen^+$ vectors, cell sorted and subcloned. WT or $KO^{CXCL5}$ clones were identified by ELISA and validated *in vitro* **B)** Representative histogram of OVA expression measured by flow cytometry in KP-OVA WT or $KO^{CXCL5}$ cells. MFI is indicated nearby the corresponding histogram. **C)** Growth curve of KP-OVA WT and $KO^{CXCL5}$ cells *in vitro*. Data represent the mean±SEM of 4 independent measurements/3 replicates each. Significance was determined by a two-way ANOVA with ns $p>0.05$. **D)** Frequencies of the indicated immune subsets in the lung of mice challenged with WT or $KO^{CXCL5}$ cells 18 days after challenge. Absolute numbers of different subsets were measured with counting beads by flow cytometry. Data represent the mean±SEM of 3-4 mice/group. Significance was determined by two-way ANOVA with ****$p\leq0.0001$. **E)** Frequencies of circulating neu were analysed at 2 time points after tumor challenge. Data represent the mean±SEM of 4 mice/group. Significance was determined by two-way ANOVA. **F)** Frequencies of neu expressed as % of total cells were analysed in mediastinal lymph node (mLN), bone marrow (BM) and spleen by flow cytometry. Data are mean±SEM of 4 mice/group. Significance was determined by unpaired t-test with **$p\leq0.01$.

**Supplementary Fig. S3. Rescue of CXCL5 expression induces neu recruitment. A)** KP-OVA $KO^{CXCL5}$ cells were transduced with CXCL5-expressing (lenti-CXCL5) or empty vector (lenti-vec). **B)** CXCL5 production by the indicated clones was assessed by ELISA. Data represent the mean±SEM of 2-9 independent measurements. Significance was determined by a one-way ANOVA with ****$p \leq 0.0001$. **C)** Relative abundance of *Cxcl5* was evaluated by qRT-PCR in lung tissues at day 9 upon challenge with the indicated KP genotypes. Data represent the mean±SEM of 3-6 independent RNA extraction. Significance was determined by one-way ANOVA with **$p \leq 0.01$, ****$p \leq 0.0001$. **D)** Representative dot plot and relative abundance of SiglecF $^{High/low}$ neu in mice challenged with KP-OVA$^{lenti\text{-}CXCL5}$ or $^{lenti\text{-}vec}$. Bars show quantification of neu frequencies on CD45$^{+}$ total lung cells and the fraction of SiglecF$^{Hi/low}$ among neu (percentages above bars refer to SiglecF$^{high}$ neu). Data represent the mean±SEM of one or two independent experiment, 3-4 mice each group.

**Supplementary Fig. S4. CD4 T cell recruitment and activation are marginally affected by neutrophils. A-B)** Mice were challenged with KP OVA WT or KO$^{CXCL5}$ tumor cells and endogenous T cell responses in mLNs were evaluated 9 days after challenge. **A)** Frequencies of activated (PD1$^{+}$LAG3$^{+}$TIM3$^{+}$), memory (CD62l$^{-}$CD44$^{+}$) and effector (EOMES$^{-}$Tbet$^{+}$) CD8$^{+}$ T cells expressed as % of total mLNs-CD8$^{+}$ T cells. Data are mean±SEM of 3-6 mice each group. Significance was determined by unpaired t-test with ns $p > 0.05$, **$p \leq 0.01$. **B)** CD8$^{+}$ T cells from mLNs of control and tumor bearing mice were *ex vivo* restimulated with SIINFEKL and production of IFN$\gamma$ was measured by intracellular staining by flow cytometry. Data represent the mean±SEM of 3 independent experiments with 2-5 mice each group. Significance was determined by one-way ANOVA with ns $p > 0.05$, *$p \leq 0.05$. **C-D)** Frequencies of activated (PD1$^{+}$LAG3$^{+}$TIM3$^{+}$) and memory (CD62L$^{-}$CD44$^{+}$) CD4$^{+}$ T cells expressed as % of total CD4$^{+}$ T cells in normal or tumor bearing lungs **C)** and corresponding mLNs **D)** are shown. Frequencies represent mean±SEM of 1 or 2 experiments with 2-4 mice/group. Significance was determined by one-way ANOVA with ns $p > 0.05$, *$p \leq 0.05$, ***$p \leq 0.001$ **C)** or by unpaired t-test with ns $p > 0.05$ **D)**.

**Supplementary Fig. S5. Low effectiveness of neu-specific antibody depletion. A-E**) Scheme of neu depletion *in vivo*. Mice were challenged with KP OVA cells and, starting from day 6$^{th}$, treated every 3 days with $\alpha$Ly6G or isotype control. **B)** Frequencies of CD11b$^{+}$Ly6G$^{+}$ (neu) in blood of challenged mice at days 7$^{th}$ and 13$^{th}$. Data are expressed as % of total blood cells and are mean±SEM of 3 mice/group. Significance was determined by two-way ANOVA with ***p≤0.001. **C)** Representative dot plot and quantification of SiglecF$^{High/Low}$-neu expressed as % of CD45$^{+}$ cells. Frequencies represent the mean±SEM of 4 mice/group. **D)** Representative sections of neu depleted tumor bearing lungs labelled with aLy6G (brown dots). Representative 10x section (left) and corresponding 40x magnification. Scale bars (100 µm) are indicated.

### Supplementary Tables

**Supplementary Table 1.** List of flow cytometry antibodies.

| Antibody | Fluorophore | Clone | Company |
|---|---|---|---|
| CD45 | APCCy7 | 30-F11 | Biolegend |
| CD3 | PerCPCy5.5 | 145-2C11 | Biolegend |
| CD3 | FITC | 145-2C11 | Biolegend |
| CD4 | BV785 | GK1.5 | Biolegend |
| CD8 | APC | 53-6.7 | Biolegend |
| CD62l | BV650 | MEL-14 | Biolegend |
| CD44 | PE | IM7 | Biolegend |
| PD-1 | EF450 | J43 | eBioscience |
| TIM3 | PerCPCy5.5 | RMT3-23 | Biolegend |
| LAG3 | BV650 | C9B7W | Biolegend |
| IFN-γ | PE | MG1.2 | Biolegend |
| EOMES | A647 | W17001A | Biolegend |
| Tbet | BV421 | 4B10 | Biolegend |
| SIINFEKL/H-2Kb Pro5 | PE | | ProImmune |
| CD11b | BV421 | M1/70 | BD Bioscience |

| | | | |
|---|---|---|---|
| SiglecF | PerCPEF710 | IRNM44N | eBioscience |
| SiglecF | BB515 | E50-2440 | BD Bioscience |
| CD11c | BV786 | N418 | Biolegend |
| CD11c | APC | N418 | Biolegend |
| MHC-II | APCR700 | M5/114.15.2 | Biolegend |
| CD86 | PE | B7-2 | BD Bioscience |
| PD-L1 | PerCPeF710 | MIH5 | eBioscience |
| CD64 | BV605 | X54-5/7.1 | Biolegend |
| Ly6C | AF488 | HK1.4 | eBioscience |
| Ly6C | BV570 | HK1.4 | Biolegend |
| Ly6G | AF488 | 1A8 | Biolegend |
| Ly6G | PE | 1A8 | Biolegend |
| CXCR2 | PerCPCy5.5 | SA044G4 | Biolegend |
| B220 | APC | C363-16A | Biolegend |
| CD19 | PE | 1D3 | BD Bioscience |
| NK1.1 | Biotin | PK136 | Biolegend |
| Streptavidin | PE-Cy7 | | Biolegend |
| L&D | BV510 | | Life Technologies |
| CD16/CD32 | | 93 | Biolegend |
| HA | | 3F10 | Roche |
| $\alpha$RAT | AF488 | | Invitrogen |

**Supplementary Methods**

**Collection and processing of human lung cancer gene expression data.**

A lung cancer compendium has been created from 7 major datasets comprising microarray data of lung cancer samples annotated with clinical outcome. All data were measured on Affymetrix arrays and have been downloaded from NCBI Gene Expression Omnibus (GEO, http://www.ncbi.nlm.nih.gov/geo/) GSE3141, GSE10245, GSE14814, GSE19188, GSE31210, GSE68465, and form the Ladanyi and Gerald Laboratories Lung Adenocarcinoma microarray repository (http://cbio.mskcc.org/public/lung_array_data/). Prior to analysis, we eliminated duplicate samples and renamed all original sets after the medical center where patients were recruited. This re-organization returned 1,136 unique samples from 10 independent cohorts comprising 989 adenocarcinomas, 778 of which with complete clinical outcome information (Table 1). The type and content of clinical and pathological annotations of the compendium samples have been derived from the original cohorts. Since raw data (.CEL files) were available for all samples, the integration, normalization and summarization of gene expression signals has been obtained applying the procedure described in Rustighi et al. (1). Briefly, expression values were generated in R from intensity signals using a custom CDF obtained merging HG-U133A, HG-U133A2 and HG-U133 Plus2 original CDFs and transforming the original CEL files accordingly. Intensity values have been background-adjusted, normalized using quantile normalization, and gene expression levels calculated using median polish summarization (multi-array average procedure, RMA). Gene expression data for a total of 21,995 probe sets have been collapsed to 12,391 unique gene symbols using the *hgu133a.db* annotation package (version 3.2.3) and the *aggregate* function of R *stats* package. Clinical information among the various datasets has been standardized as described in Cordenonsi et al. (2). Average signature expression has been calculated as the average expression of all signature genes in sample subgroups.

**Human neutrophil fraction analysis**

Neutrophil cell fractions have been quantified using CIBERSORT (3)on the lung adenocarcinoma samples of the *LUAD compendium*. Briefly, the non-log linear expression matrix with the 989 cases was uploaded to the *CIBERSORT* R script (version 1.04) as a mixture file and *CIBERSORT* was run in absolute mode with the LM22 signature gene file, 100 permutations, and quantile normalization. In absolute mode, CIBERSORT scales relative cellular fractions into a score that reflects the absolute proportion of each cell type in a mixture. Although not expressed as a fraction, the absolute score can be directly compared both between- and within-samples (4). Samples have been divided into three groups based on the upper (high neutrophil content) and lower (low neutrophil content) quartiles of the absolute scores for the neutrophil cell type. Samples that fell in between the quartile cut offs have been termed the medium neutrophil content group (5).

**Survival analysis**

To evaluate the prognostic value of the neutrophil content, we estimated the overall survival probability of the high and low content groups using the Kaplan–Meier method and compared the Kaplan–Meier curves using the log-rank (Mantel–Cox) test. P values were calculated according to the standard normal asymptotic distribution. To evaluate the prognostic level of the neutrophil signature from Zilionis et al. (6) we separated the samples into two groups based on a signature score obtained summarizing the standardized expression levels of the signature genes into a combined score with zero mean (7). Tumor samples have been classified as signature 'Low' if the combined score was negative and as signature 'High' if the combined score was positive. The overall survival probability in the two groups has been estimated using the Kaplan–Meier method and the Kaplan–Meier curves compared using the log-rank (Mantel–Cox) test. P values were calculated according to the standard normal asymptotic distribution. Survival analysis was performed in GraphPad Prism.

1. Rustighi A, Zannini A, Tiberi L, Sommaggio R, Piazza S, Sorrentino G, et al. Prolyl-isomerase Pin1 controls normal and cancer stem cells of the breast. EMBO Mol Med. 2014;6(1):99-119.
2. Cordenonsi M, Zanconato F, Azzolin L, Forcato M, Rosato A, Frasson C, et al. The Hippo transducer TAZ confers cancer stem cell-related traits on breast cancer cells. Cell. 2011;147(4):759-72.
3. Newman AM, Liu CL, Green MR, Gentles AJ, Feng W, Xu Y, et al. Robust enumeration of cell subsets from tissue expression profiles. Nat Methods. 2015;12(5):453-7.
4. Sturm G, Finotello F, Petitprez F, Zhang JD, Baumbach J, Fridman WH, et al. Comprehensive evaluation of transcriptome-based cell-type quantification methods for immuno-oncology. Bioinformatics. 2019;35(14):i436-i45.
5. Craven KE, Gokmen-Polar Y, Badve SS. CIBERSORT analysis of TCGA and METABRIC identifies subgroups with better outcomes in triple negative breast cancer. Sci Rep. 2021;11(1):4691.
6. Zilionis R, Engblom C, Pfirschke C, Savova V, Zemmour D, Saatcioglu HD, et al. Single-Cell Transcriptomics of Human and Mouse Lung Cancers Reveals Conserved Myeloid Populations across Individuals and Species. Immunity. 2019;50(5):1317-34 e10.
7. Adorno M, Cordenonsi M, Montagner M, Dupont S, Wong C, Hann B, et al. A Mutant-p53/Smad complex opposes p63 to empower TGFbeta-induced metastasis. Cell. 2009;137(1):87-98.
